# Subcutaneous nanotherapy repurposes the immunosuppressive mechanism of rapamycin to enhance allogeneic islet graft viability

**DOI:** 10.1101/2020.09.03.281923

**Authors:** Jacqueline A. Burke, Xiaomin Zhang, Sharan Kumar Reddy Bobbala, Molly A. Frey, Carolina Bohorquez Fuentes, Helena Freire Haddad, Sean D. Allen, Reese A.K. Richardson, Guillermo A. Ameer, Evan A. Scott

## Abstract

Oral rapamycin administration rapamycin is plagued by poor bioavailability and wide biodistribution. Thus, this pleotropic mTOR inhibitor has a narrow therapeutic window, numerous side effects and provides inadequate transplantation protection. Parental formulation was not possible due to rapamycin’s hydrophobicity (log P 4.3). Here, we demonstrate that subcutaneous rapamycin delivery via poly(ethylene glycol)-*b*-poly(propylene sulfide)(PEG-*b*-PPS) polymersome (PS) nanocarriers modulates cellular biodistribution of rapamycin to change its immunosuppressive mechanism for enhanced efficacy while minimizing side effects. While oral rapamycin inhibits naïve T cell proliferation directly, subcutaneously administered rapamycin-loaded polymersomes (rPS) instead modulated Ly-6C^low^ monocytes and tolerogenic semi-mature dendritic cells, with immunosuppression mediated by CD8+ Tregs and rare CD4+ CD8+ double-positive T cells. As PEG-*b*-PPS PS are uniquely non-inflammatory, background immunostimulation from the vehicle was avoided, allowing immunomodulation to be primarily attributed to rapamycin’s cellular biodistribution. Repurposing mTOR inhibition significantly improved maintenance of normoglycemia in a clinically relevant, MHC-mismatched, allogeneic, intraportal (liver) islet transplantation model. These results demonstrate the ability of engineered nanocarriers to repurpose drugs for alternate routes of administration by rationally controlling cellular biodistribution.

## Introduction

Type 1 diabetes (**T1D**) is an endocrine disorder that leads to pancreatic β cell destruction and requires management with lifelong exogenous insulin therapy^1^. With the advent of the Edmonton protocol, islet transplantation has emerged as a promising treatment for T1D by eliminating the need for exogenous insulin^1^. This protocol involves three key components: acquisition of viable insulin-producing cells, surgical transplantation of these cells into a suitable physiological location to maintain glucose sensitivity and responsiveness, and an immunosuppressive regimen to maintain islet viability and protection from the host immune system^1^. While all three components remain active areas of research, the need for immunosuppression remains the key limitation preventing islet transplantation from becoming the clinical standard of care for all T1D patients^1,2^. A critical advancement in this regard was the advent of orally administered nanocrystal rapamycin, i.e Rapamune, for non-steroidal immunosuppressive protocols. Rapamycin inhibits the mammalian target of rapamycin (**mTOR**) pathway to directly inhibit T cell proliferation by arresting these cells in the G1 phase of the cell cycle and preventing IL-2 secretion^3^. Although more effective than prior immunosuppressive protocols involving intravenous steroid administration, patients undergoing transplantation procedures are still plagued by frequent graft rejection and an unpleasant array of side effects^3,4^.

Side effects related to oral rapamycin administration stem primarily from poor and inconsistent bioavailability and the wide cellular biodistribution. Rapamune has a bioavailability of only 14% in the solution form and 41% in tablet form^3^. The low bioavailability is attributed primarily to the first pass metabolism associated with the oral route of administration, cytochrome P450 elimination and transport by p-glycoprotein efflux pumps. For example, absorption of Rapamune is greatly varied by fat content in food, and cytochrome P450 isoenzyme CYP3A4 metabolism can cause serious drug-drug interactions^3^. With regards to biodistribution, lipophilic Rapamune primarily partitions into red blood cells (95%) and then eventually accumulates in off-target organs, including the heart, kidneys, intestines and testes^5-7^, leading to side effects. These side effect occur due to the ubiquitous expression of mTOR in diverse cell types, resulting in unintended cell populations also experiencing cell cycle arrest^2,3^. Clinically, this can lead to malignancy, enhanced susceptibility to infection, impaired wound healing, thrombopenia, alopecia, gastrointestinal distress, gonadal dysfunction, hypertension, hyperlipidemia, nephrotoxicity and peripheral edema^3,8^. In order to balance the need to maintain immunosuppression with the avoidance of side effects, patients must undergo frequent blood work to ensure that the rapamycin concentration is within the small therapeutic window of 5 to 15 ng/ml^3,5^. Of note, mTOR inhibition can have distinct responses depending on the cell type. For example, rapamycin retains dendritic cells (**DCs**) in an immature tolerogenic state that resists coreceptor expression in response to inflammatory stimuli, a process known as costimulation blockade^9^.

Given the plethora of problems associated with oral Rapamune, an alternative therapy that bypasses the oral route of administration, reduces adverse effects, and enhances transplantation outcomes is needed. Subcutaneous administration would avoid bioavailability issues that plague Rapamune including first pass metabolism, elimination by intestinal cytochrome P450 and p-glycoprotein, and variability associated with food content^3^. Importantly, the subcutaneous route of administration provides the advantage of targeting lymphatic drainage^10^. Unlike intravenous administration, the subcutaneous route would allow patients to take their medication from their own home. The T1D patient population is well versed in the subcutaneous method of injection due to the need to inject insulin. However, due to the lipophilic nature of rapamycin (log P 4.3), it is poorly soluble and therefore very difficult to formulate into a parental drug for subcutaneous administration^11^. Additionally, subcutaneous administration of rapamycin is unlikely to achieve the required biodistribution to directly inhibit T cell proliferation and prevent islet rejection. But the subcutaneous route does provide access to antigen presenting cells (**APCs**), including the aforementioned DCs that can elicit potent tolerogenic responses upon modulation by rapamycin. Tolerogenic DCs (**tDCs**) constitutively generate regulatory T cells (**Tregs**) as well as express anti-inflammatory cytokines, both of which have been linked to enhanced survival of transplanted islets^10^.

We hypothesized that focusing rapamycin’s mTOR inhibition on APCs using engineered nanocarriers could achieve sustained immunosuppression and survival of transplanted islets via the subcutaneous route with lower dosage and minimal side effects (**Fig. 1**). To focus the broad biodistribution of rapamycin specifically to APCs, we generated rapamycin-loaded poly(ethylene glycol)-*b*-poly(propylene sulfide)) (**PEG-*b*-PPS**) polymersomes (**rPS**). The PEG-*b*-PPS polymersome (**PS**) platform allows for efficient loading of lipophilic drugs within the PPS membrane^12^, has been validated to be nontoxic in both mice and nonhuman primates^13-17^ and undergoes uptake by DC and monocyte populations^17^, which are critically responsible for directing T cell activation during immune responses^15,18,19^. Importantly, PEG-*b*-PPS is non-immunomodulatory relative to other common nanomaterials, with an immunostimulatory profile that is determined almost exclusively by the loaded therapeutic^17^. Unloaded PEG-*b*-PPS PS elicit minimal immunomodulatory activity, whereas comparable poly(lactic-co-glycolic acid) (**PLGA**) nanocarriers cause an extensive immunomodulatory response, including alteration of immune cell populations, changes in coreceptor expression (e.g. CD80, CD86), and modification of the inflammatory status of numerous immune cell subsets^20^. Thus, our mechanistic assessment of rPS-mediated immunosuppression avoids interference from background immunomodulation by the drug vehicle itself.

**Fig. 1.**
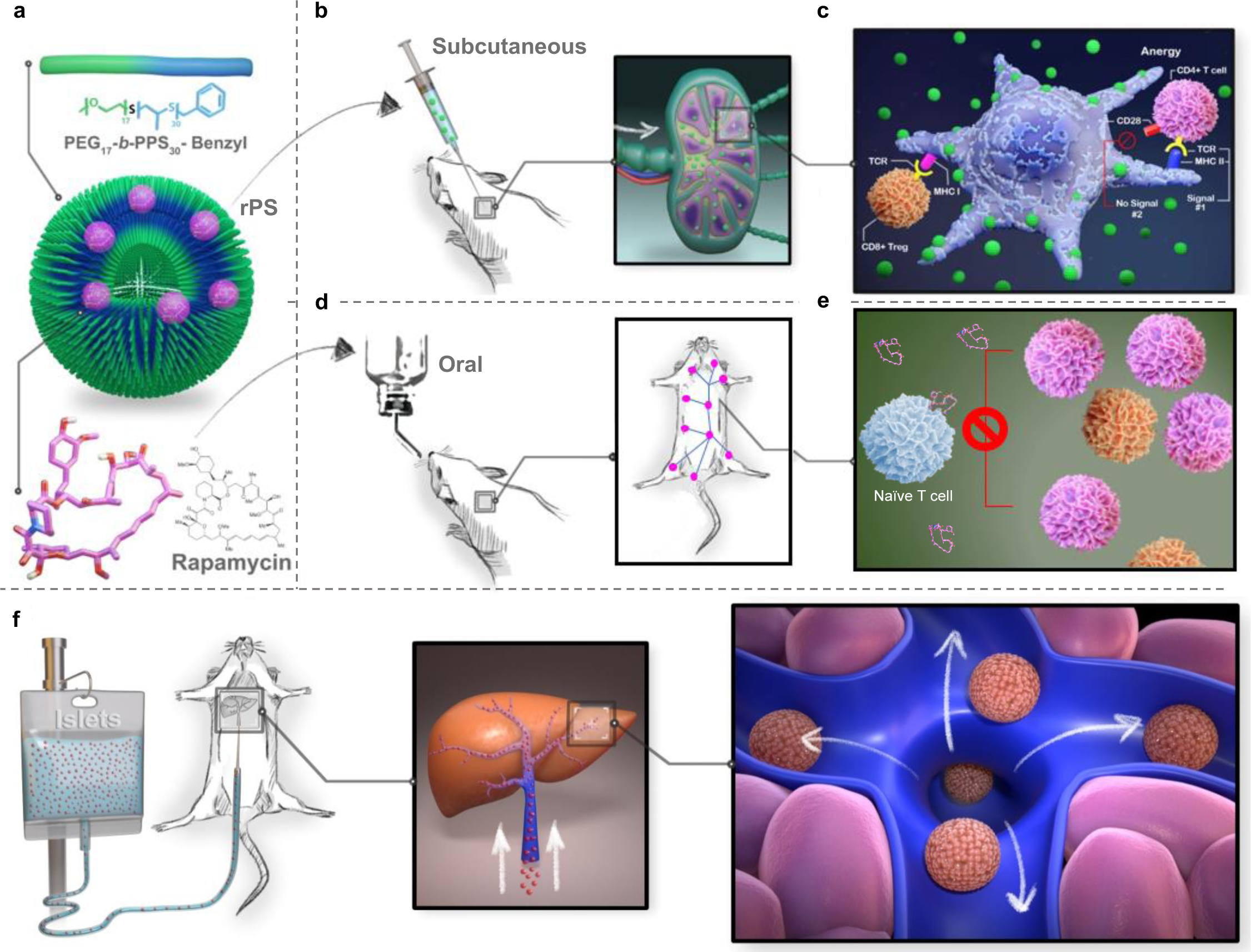
Subcutaneous rapamycin delivery to APCs via rPS signals T cells to tolerate fully-MHC mismatched intraportal islet graft. **a**, Rapamycin can be easily loaded into the hydrophobic membrane of polymersomes (PS) to form rapamycin-loaded polymersomes (rPS). **b**, When injected subcutaneously into mice, rPS drain into the brachial lymph nodes where they are uptaken by antigen-presenting cells (APCs). **c**, As a result, APCs develop an anti-inflammatory, semi-mature phenotype, in which they express high levels of MHC II to present to CD4+ T cell receptors, but they do not express costimulatory molecules. Without activation from costimulation, acute rejection-causing CD4+ T cells go into a state of anergy or become tolerogenic CD8+ regulatory T cells (Tregs). **d**, Clinically, rapamycin is given orally. Oral administration results in a wide biodistribution and low bioavailability. **e**, Oral rapamycin primarily acts on T cells directly by inhibiting mTOR and thus preventing T cell proliferation. **f**, To assess the ability of subcutaneous rPS to provide a tolerogenic state that allows for fully-major histocompatibility complex (MHC) mismatched allogeneic graft survival, islet transplantation was performed in diabetic mice at the clinically relevant intraportal (liver) transplantation site and graft viability was assessed by the restoration and maintenance of normoglycemia.

To the best of our knowledge, herein we present the first therapeutic application of subcutaneous rapamycin nanotherapy, as well as the first reorchestrating of the immunological mechanism of an immunosuppressant by rationally controlling its cellular biodistribution. Efficacy of this strategy is assessed via high parameter spectral flow cytometry with analysis via T-distributed stochastic neighbor embedding (**tSNE**), single-cell RNA sequencing and a clinically relevant intraportal fully-major histocompatibility complex (**MHC**) mismatched allogeneic islet transplantation model. Our results reveal the mechanisms behind how nanocarrier-mediated modulation of cellular biodistribution can significantly change the therapeutic window, reduce adverse events, and enhance anti-inflammatory efficacy of an immunosuppressant by rationally changing its therapeutic mechanism of action.

## Results

### Rapamycin loading does not alter PS morphology or polydispersity

PEG-*b*-PPS PS were characterized to assess encapsulation efficiency and retention of their vesicular nanostructure following the loading of rapamycin to form rPS. Rapamycin encapsulation efficiency was found to be greater than 55% for rPS following self-assembly and therapeutic loading via thin film hydration of desiccated PEG-*b*-PPS films. Neither the PS vesicular nanostructure nor the polydispersity were significantly modulated by rapamycin loading as assessed by dynamic light scattering (**DLS**), cryogenic transmission electron micrograph (**cryoTEM**) and small angle x-ray scattering (**SAXS**) (**Fig. 2a-c**). The stability of rapamycin loading was assessed in 1X phosphate buffered saline (**PBS**) at 4 °C, finding approximately 94% of the drug was retained over the course of 1 month (**Fig. S1**). Rapamycin is relatively lipophilic with a logP of 4.3^11^, and thus these results were consistent with past attempts to load molecules of low water solubility into PEG-*b*-PPS nanostructures.

**Fig. 2.**
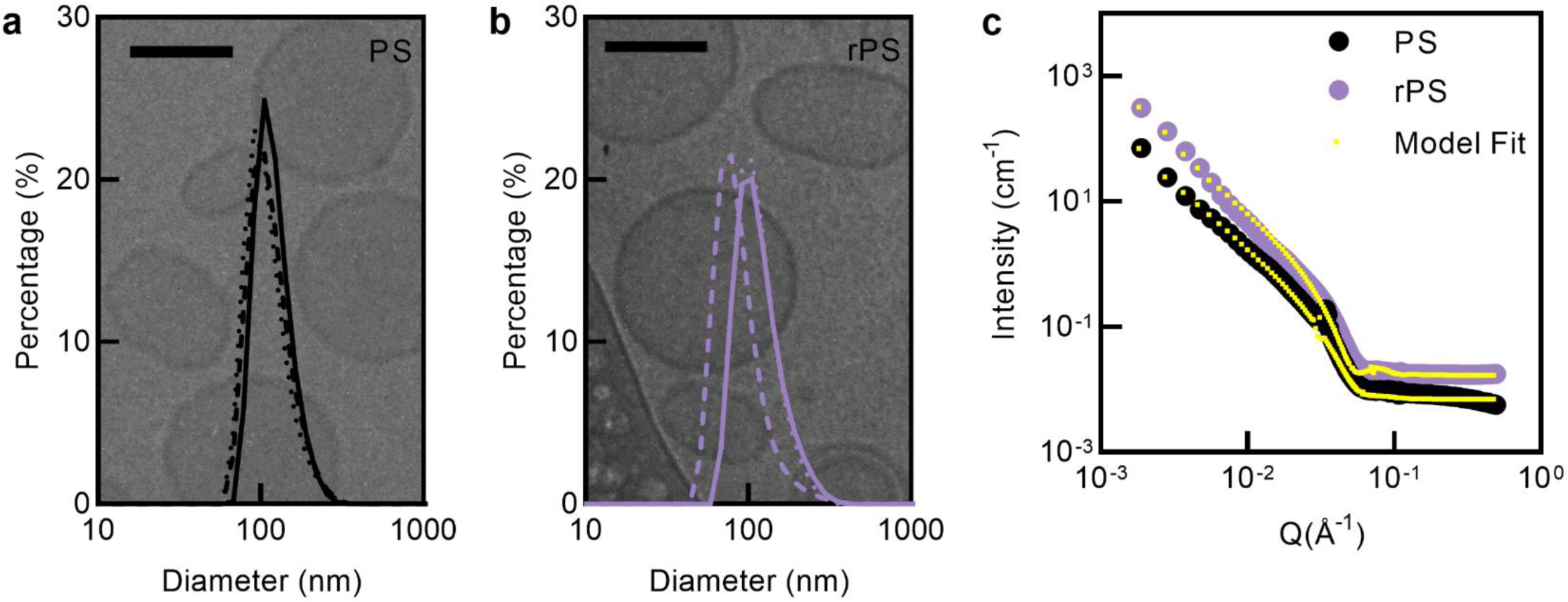
Morphological characterization of PS and rPS. **a,b**, Cryogenic transmission electron micrograph (cryoTEM) of polymersomes (PS) (**a**) and rapamycin-loaded polymersomes (rPS) (**b**) with overlay of size distribution by dynamic light scattering (DLS) (n = 3). Scale bars represent 100 nm. (n = 3). **c**, Small angle x-ray scattering (SAXS) transformed data of PS (●) and rPS 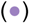 with polymer vesicular model fit 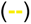. (n = 3-5).

### Subcutaneous rapamycin delivery via PS alters biodistribution and immunomodulation

To demonstrate that PEG-*b*-PPS PS can alter the biodistribution of a small molecule following subcutaneous administration, indocyanine green dye (**ICG-PS**) was loaded into PS to serve as a traceable model payload. We subcutaneously injected C57BL/6J mice with ICG-PS or free ICG, sacrificed animals at 2, 24, and 48 h post injection and analyzed organs via IVIS (**Fig. S2**). We show that ICG-PS allowed for sustained residence within to the superficial axillary/brachial lymph nodes at 24 and 48 h post-injection, whereas free form ICG dye had been cleared at these later time points (**Fig. 3a**). To confirm that this effect holds true for rapamycin, rapamycin (in 0.2% carboxymethyl cellulose (**CMC**)) or rPS were subcutaneously injected into C57BL/6J mice and the animals were sacrificed at 0.5, 2, 8, 18, 24, and 48 h post-injection to assess rapamycin content in various organs. We found that delivery of rapamycin via rPS increases rapamycin concentration in immune cell-rich tissues, such as the blood, liver, axillary lymph center (deep axillary/axillary/axial and superficial axillary/brachial lymph nodes; **AX LN**), subiliac lymph center (subiliac/inguinal lymph nodes; **IN LN**) and spleen (**Fig. 3b**). To assess both the organ and cellular effects of rapamycin delivery via PEG-*b*-PPS PS, immune cell populations from various tissues were isolated after repeated subcutaneous injection with unloaded PS, rapamycin in CMC or rPS formulations (**Fig. 3c**). When unloaded PS were injected, very little immunomodulation was observed (**Fig. 3c**). However, when rapamycin was loaded with in the PS, potent immunomodulation occurred (**Fig. 3c**). The relative immunologically inert status of the unloaded PS allowed for the majority of the effects of rPS to be attributed to the altered biodistribution of the drug, as opposed to the nanocarrier itself. A significant change in immunomodulation is observed for PS-mediated rapamycin delivery (**Fig. 3c**). Taken in combination with the inert nature of PEG-*b*-PPS, our results demonstrate that rapamycin’s organ and cellular biodistribution have a strong influence on the resulting immunological effect.

**Fig. 3.**
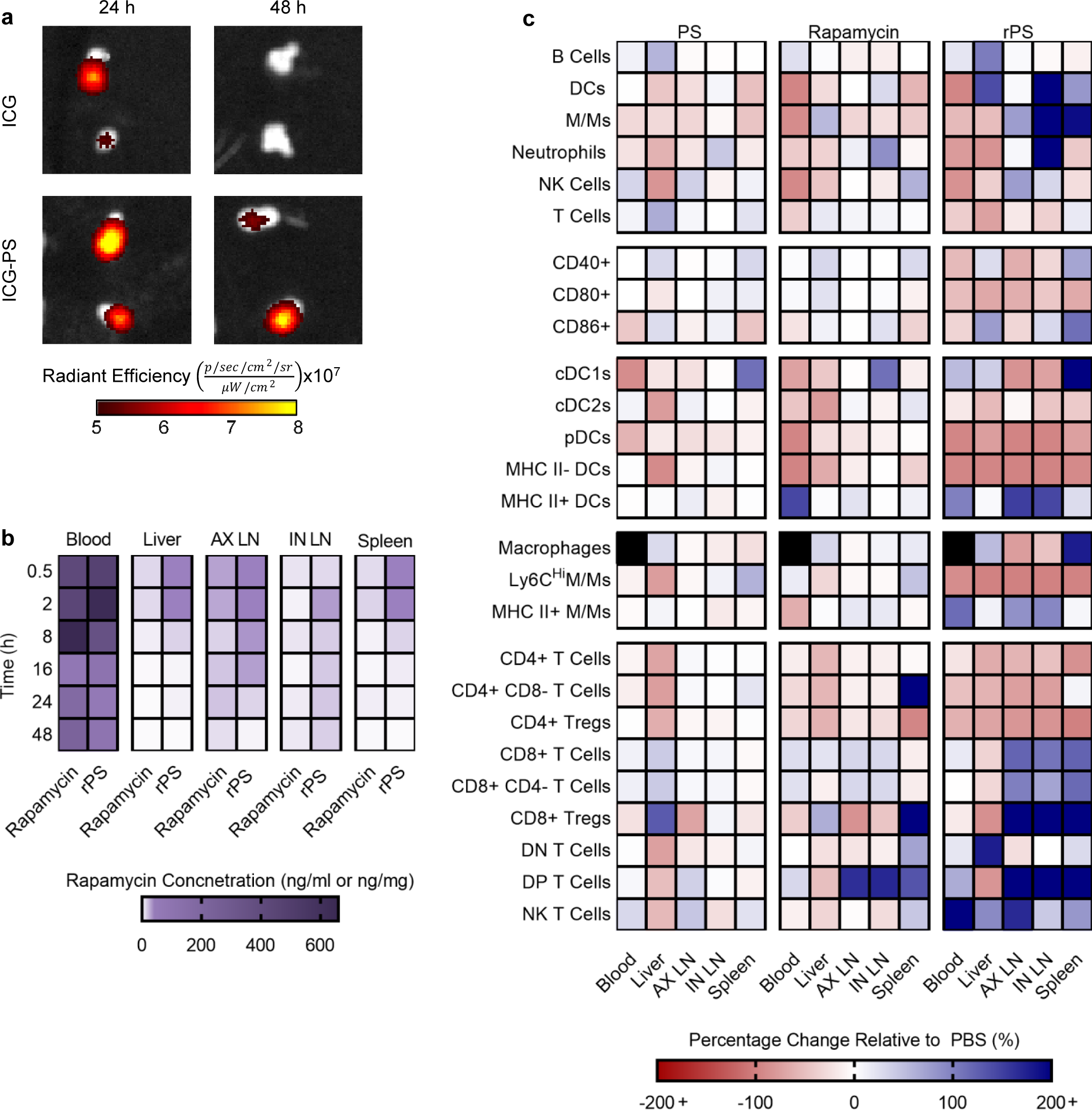
Subcutaneous rapamycin delivery via PS alters biodistribution and immunomodulation. **a**, Biodistribution of indocyanine green (ICG) dye in the superficial axillary/brachial LN (of the axillary lymphocenter (AX LN)) 24 and 28 hours after subcutaneous injection (n = 5 mice/group) with ICG or ICG-loaded polymersomes (ICG-PS). **b**, Biodistribution of rapamycin after subcutaneous injection with rapamycin or rapamycin-loaded polymersomes (rPS). Rapamycin concentration in the blood, liver, axillary lymphocenter (deep axillary/axillary/axial and superficial axillary/brachial lymph nodes; AX LN), subiliac lymphocenter (subiliac/inguinal lymph nodes; IN LN) and spleen over time (0.5 h, 2 h, 8 h, 16 h, 24 h and 48 h) as assessed by LC-MS/MS. (n = 3 mice/group). **c** Flow cytometry analysis of CD45+ cell populations from mice subcutaneously injected with 1X phosphate buffered saline (PBS), polymersomes (PS), rapamycin, or rPS for 11 days at a dose of 1 mg/kg/day rapamycin or equivalent. Macrophages were not assessed in blood as indicated by a black box. All data is presented as mean percentage change relative to PBS treatment. (n = 3 mice/group).

### Costimulation blockade and MHC II upregulation via rPS induces CD4+ T cell depletion and anergy

To more deeply characterize changes in immune cell populations in response to rPS delivery, we subcutaneously injected healthy mice with PBS, unloaded blank PS, rapamycin, or rPS (11 injections, 1 mg/kg rapamycin or equivalent) and subsequently extracted a variety of organs (blood, liver, AX LN, IN LN, and spleen) for assessment via high-parameter spectral flow cytometry. tSNE with Barnes-Hut approximations was used to visualize the data after gating (**Fig. S4**), down sampling, and concatenation of treatment groups. To further understand the changes in immune cell populations as a result of rPS treatment, the inflammatory state of APC populations was assessed via receptor expression. Specifically, CD40, CD80 and CD86 coreceptor presentation on B cells (**Fig. S5-9**), DCs (**Fig. 4a-c, S5a-9a**) and monocyte/macrophage linage cells (**M/Ms**)^21^ (**Fig 4e-g, S5-9**) was analyzed. Furthermore, MHC II presentation was assessed on DCs and M/Ms (**Fig 4d,h S5b-9b**). With rPS, costimulation blockade is observed as indicated by the significant downregulation of CD40, CD80, and CD86 (**Fig. 4a-c,e-g, S5a-9a**)^22^. Furthermore, rPS enhances MHC II+ APCs (**Fig 4d,h S5b-9b**). Opposing expression by MHC and coreceptors causes anergy and depletion of the CD4+ T cell population (**Fig 4i-m, S10-14**). With rPS treatment, the overall T cell population is significantly reduced (**Fig 4i, S10-14**). Further analysis reveals that this overall reduction is indeed caused by a significant decrease in CD4+ T cells and CD4+ Treg counter parts (**Fig 4j,k, S10-14**). Any remaining CD4+ T cells are left in a state of anergy as indicated by the significant decrease in CD3 and CD4 expression (**Fig 4l,m, S10-14**). These effects are most potent in the AX LN near the site of subcutaneous injection, but also occur to various lesser extents in blood, liver, IN LN, and spleen (**Fig 4, S10-14**).

**Fig. 4.**
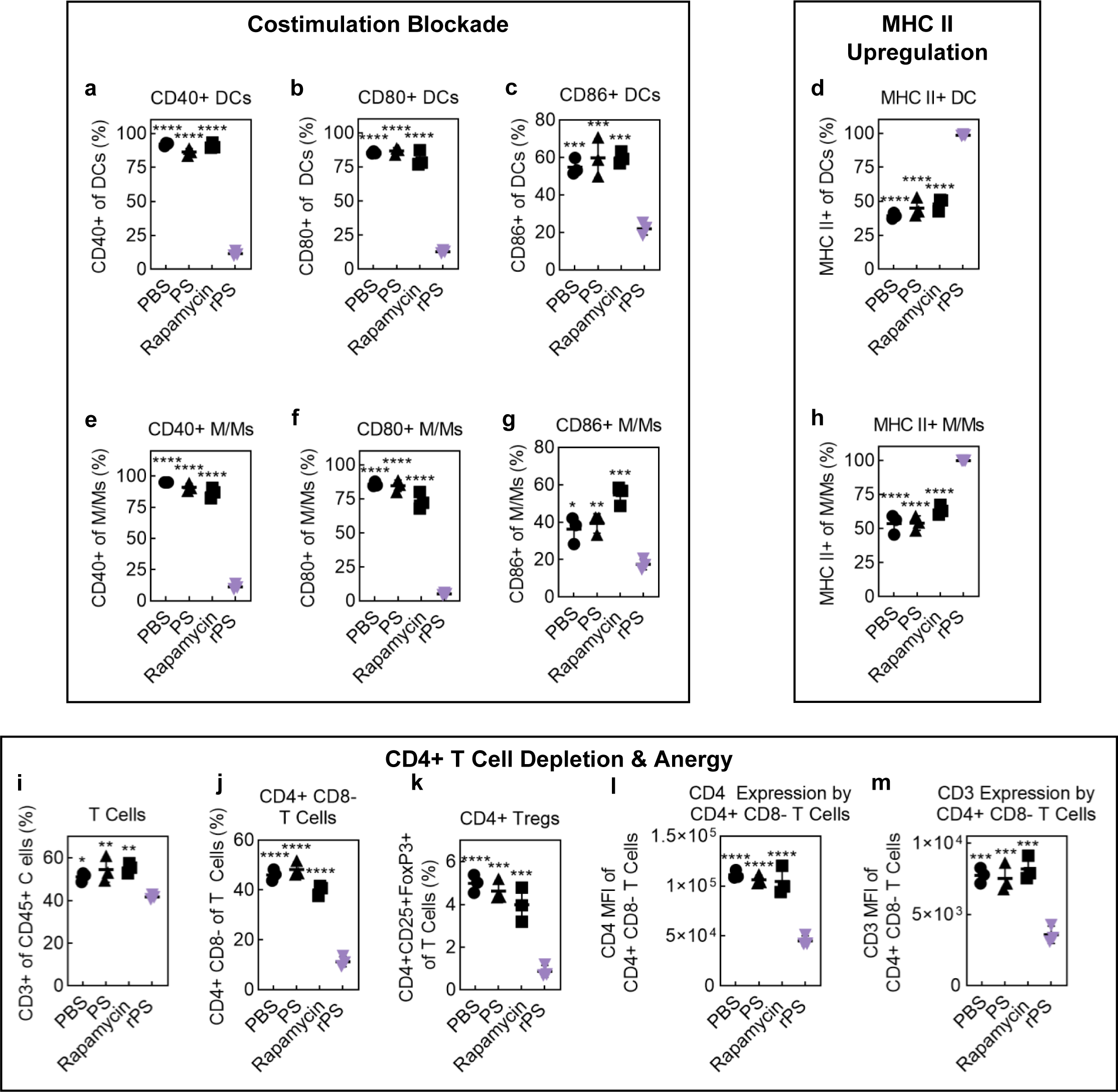
Costimulation blockade and MHC II upregulation via rPS induces CD4+ T cell depletion and anergy. Flow cytometry analysis of CD45+ cell populations from mice subcutaneously injected with 1X phosphate buffered saline (PBS;●), polymersomes (PS; ▴), rapamycin (▪) or rapamycin-loaded polymersomes (rPS; 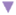) using the standard dosage protocol (11 injections, 1 mg/kg rapamycin or equivalent). **a-h)** Analysis of costimulation and major-histocompatibility complex (MHC) II: percentage of CD40+ (**a,e**), CD80+ (**b,f**), CD86+ (**c,g**) and MHC II+ (**d,h**) dendritic cells (DCs) (**a-d**) and monocyte-and-macrophage-linage cells (M/Ms) (**e-h**). **i-m**) Analysis of T cells: percentage of T cells (of CD45+ cells) (**i**), percentage of CD4+ CD8-T cells (of T cells) (**j**), percentage of CD4+ Tregs (of T cells) (**k**), CD4 expression by CD4+ CD8-T cells (MFI) (**l**) and CD3 expression by CD4+ CD8-T cells (MFI) (**m**). Data presented above is from the axillary lymphocenter (deep axillary/axillary/axial and superficial axillary/brachial lymph nodes; AX LN),. Data from the blood, liver, subiliac lymphocenter (subiliac/inguinal lymph nodes; IN LN), and spleen is shown in the supplemental Fig. S4-S14. All data are presented as mean percentage or median florescent intensity (MFI) ± SD with *p<0.05; ** p<0.01; *** p<0.001; **** p<0.0001 relative to rPS treatment. Statistical significance was determined by one-way ANOVA with Tukey’s multiple comparisons test. (n = 3 mice/group).

### rPS induce regulatory cross-talk between DCs and CD8+ T cells in lymph nodes

rPS treatment causes a significant increase in DCs within AX LN and IN LN (**Fig 5a**). More specifically, an increase in novel CD8+ CD11b+ double positive (DP) conventional DCs (cDCs) is observed (**Fig 5b**). Despite the overall increase in the DC population, plasmacytoid DC (pDC) are significantly reduced (**Fig 5c**). A significant decrease in the overall T cell population was observed due to a significant decrease in CD4+ T cells **(Fig. 4i,j**). As a result, CD8+ T cells take over a larger portion of the T cell population (**Fig 5d**), accompanied by a significant upregulation of CD8+ Tregs (**Fig 5e**) and CD8+ NK T cells (**Fig 5f,g**).

**Fig. 5.**
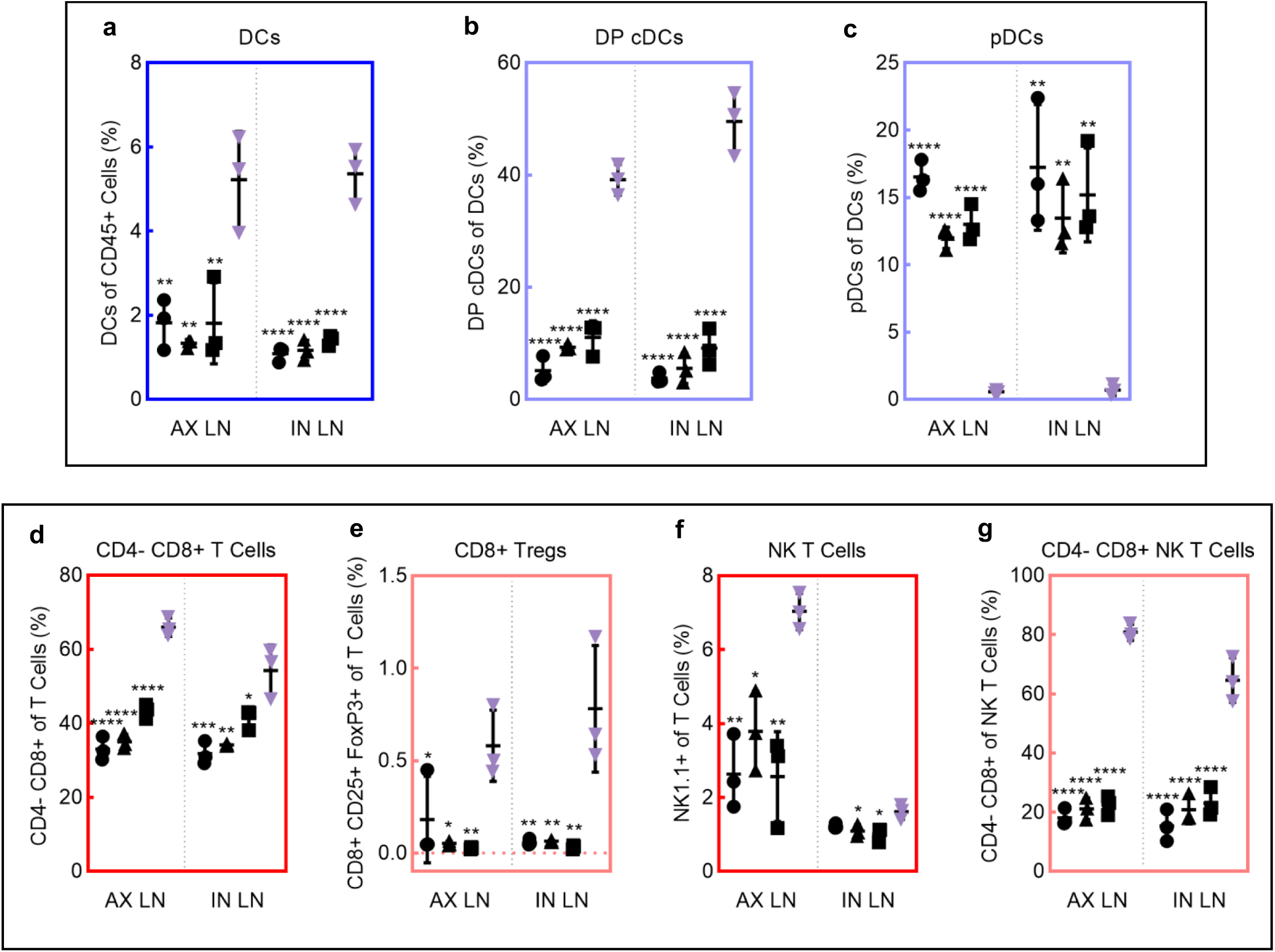
rPS induce regulatory cross-talk between DCs and CD8+ T cells in lymph nodes. Flow cytometry analysis of CD45+ cell populations from the axillary lymphocenter (deep axillary/axillary/axial and superficial axillary/brachial lymph nodes; AX LN) and subiliac lymphocenter (subiliac/inguinal lymph nodes; IN LN) of mice subcutaneously injected with 1X phosphate buffered saline (PBS; ●), polymersomes (PS; ▴), rapamycin (▪) or rapamycin-loaded polymersomes (rPS; 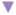) using the standard dosage protocol (11 injections, 1 mg/kg rapamycin or equivalent). **a-c**) Analysis of dendritic cells (DCs): percentage of DCs (**a**) of CD45+ cells, percentage of pDCs of DCs (**b**), percentage of DP cDCs of DCs (**c**). **d-g**) Analysis of CD8+ T cell populations: percentage of CD4-CD8+ T cells of T cells (**d**), percentage of CD8+ Tregs of T cells (**e**), percentage of NK T cells of T cells (**f**), percentage of CD4-CD8+ NK T Cells of NK T cells (**g**). All data are presented as mean percentage ± SD with *p<0.05; ** p<0.01; *** p<0.001; **** p<0.0001 relative to rPS treatment. Statistical significance was determined by one-way ANOVA with Tukey’s multiple comparisons test. (n = 3 mice/group).

### rPS induce tissue-specific suppressive Ly-6C^Lo^ M/Ms phenotypes

rPS treatment causes a significant upregulation of M/Ms in the blood, AX LN, IN LN and the spleen (**Fig. 6a**). These M/Ms are predominantly Ly-6C^Lo^ (**Fig. 6b**). In the lymph nodes, the M/Ms are most commonly monocytes, but in the spleen, macrophages dominate (**Fig. 6c**). To further understand the specific nature of M/M immunomodulation with rPS treatment, phenotypic analysis of the M/M population was performed on each tissue. Consideration for CD40, CD80, CD86, MHC II, Ly-6C and macrophage markers (F4/80 and/or CD169) was used to assign cells to one of 32 phenotypes for blood (no macrophage markers) or 64 phenotypes for LN and spleen. Phenotypic analysis reveals that rPS treatment promotes the dominance of a single suppressor M/M phenotype for each tissue, while rapamycin and control treatments present a diverse range of M/M phenotypes with often contradicting inflammatory statuses. In blood, control treatments result in a majority of CD40+ CD80+ CD86-Ly6-C^Hi^ MHC II-monocytes, while rPS treatment pushes M/Ms towards a CD40-CD80+ CD86-Ly6-C^Lo^ MHC II+ phenotype (**Fig. 6d**). rPS treated LNs are predominantly CD40-CD80-CD86-Ly6-C^Lo^ MHC II+ monocytes (**Fig. 6e,d**). rPS treated spleens are predominantly CD40+ CD80-CD86+ Ly6-C^Lo^ MHC II+ macrophages (**Fig. 6d**).

**Fig. 6.**
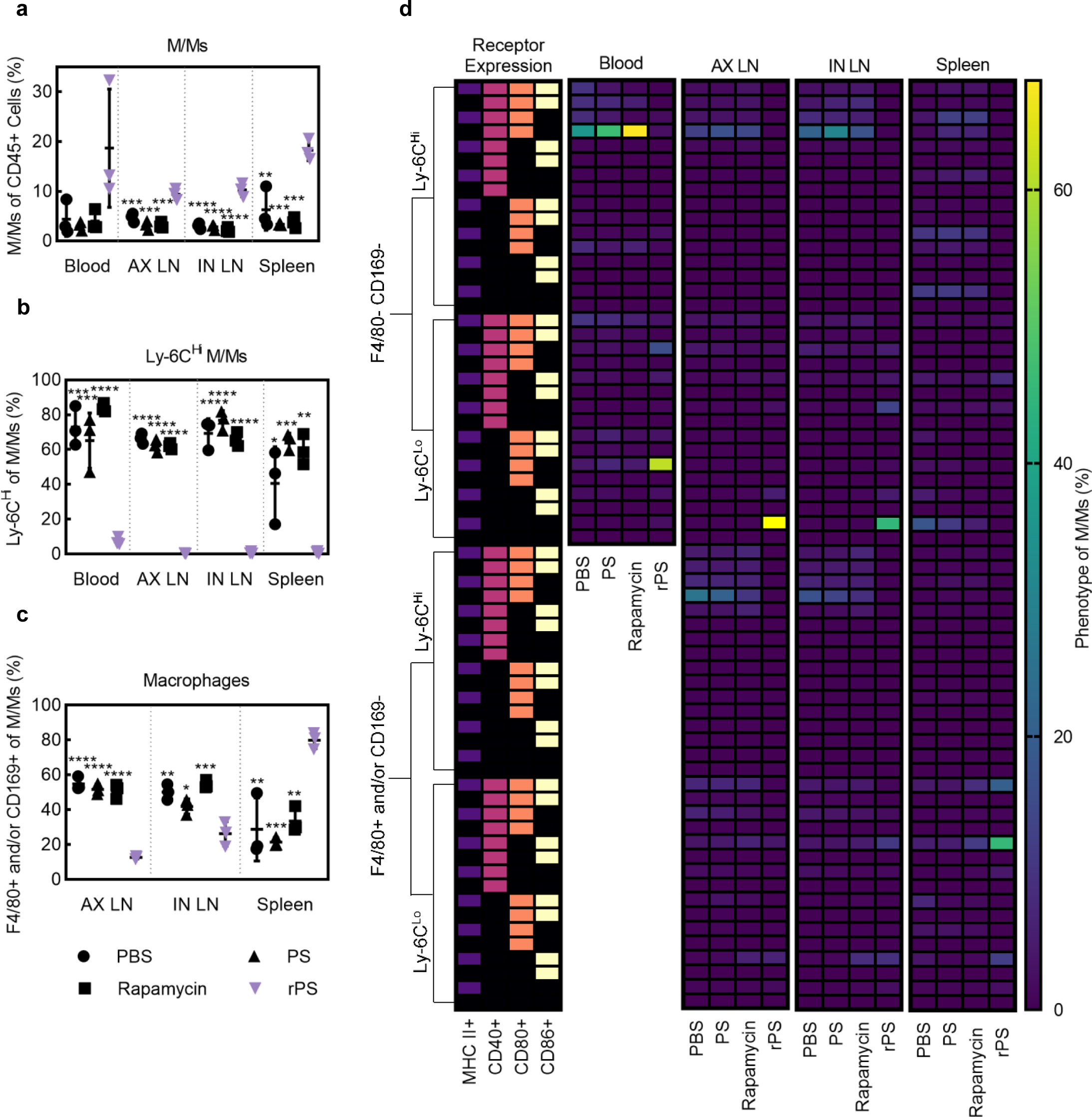
rPS induce tissue-specific suppressive Ly-6C^Lo^ M/Ms phenotypes. Flow cytometry analysis of CD45+ cell populations from mice subcutaneously injected with 1X phosphate buffered saline (PBS;●), polymersomes (PS; ▴), rapamycin (▪) or rapamycin-loaded polymersomes (rPS; 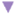) using the standard dosage protocol (11 injections, 1 mg/kg rapamycin or equivalent). **a-c**) Analysis of monocyte-and-macrophage-linage (M/M) populations in blood, axillary lymphocenter (deep axillary/axillary/axial and superficial axillary/brachial lymph nodes; AX LN), subiliac lymphocenter (subiliac/inguinal lymph nodes; IN LN) and spleen: percentage of M/Ms of CD45+ cells (**a**), percentage of Ly-6C^Hi^ M/Ms of M/Ms (**b**), percentage of macrophages (F4/80+ and/or CD169+) of M/Ms (**c**). All data are presented as mean percentage ± SD with *p<0.05; ** p<0.01; *** p<0.001; **** p<0.0001 relative to rPS treatment. Statistical significance was determined by one-way ANOVA with Tukey’s multiple comparisons test. **d**) Analysis of M/M populations by phenotype with consideration for macrophage markers (F4/80 and/or CD169; with the except of blood where macrophages were not considered), Ly-6C, MHC II, CD40, CD80, and CD86. Data are presented as heatmaps displaying the percentage of the overall M/M population occupied by each of the 32 phenotypes in the blood and 64 phenotypes in the AX LN, IN LN, and spleen. (n = 3 mice/group).

### rPS induce DP CD4^bright^ CD8^dim^ T cells with suppressor functions in lymph nodes

Interestingly, with rPS treatment, DP CD4+ CD8+ T cells have significantly larger populations in the AX LN and IN LN (**Fig. 7a,d**). From, tSNE visualization, it is observed that DP and CD4+ CD8-T cells cluster together (**Fig. 7b**). Using expression level analysis, CD4 expression by DP T cells in the rPS treatment group is similar to that of CD4+ CD8-T cells, whereas CD8 expression by DP T cells is significantly reduced as compared to CD4-CD8+ T cells. The relationship between the expression of the DP and single positive (SP) T cells can be quantified as a DP:SP expression ratio. Thus, the rPS treated DP T cell population is deemed CD4^bright^ CD8^dim^.

**Fig. 7.**
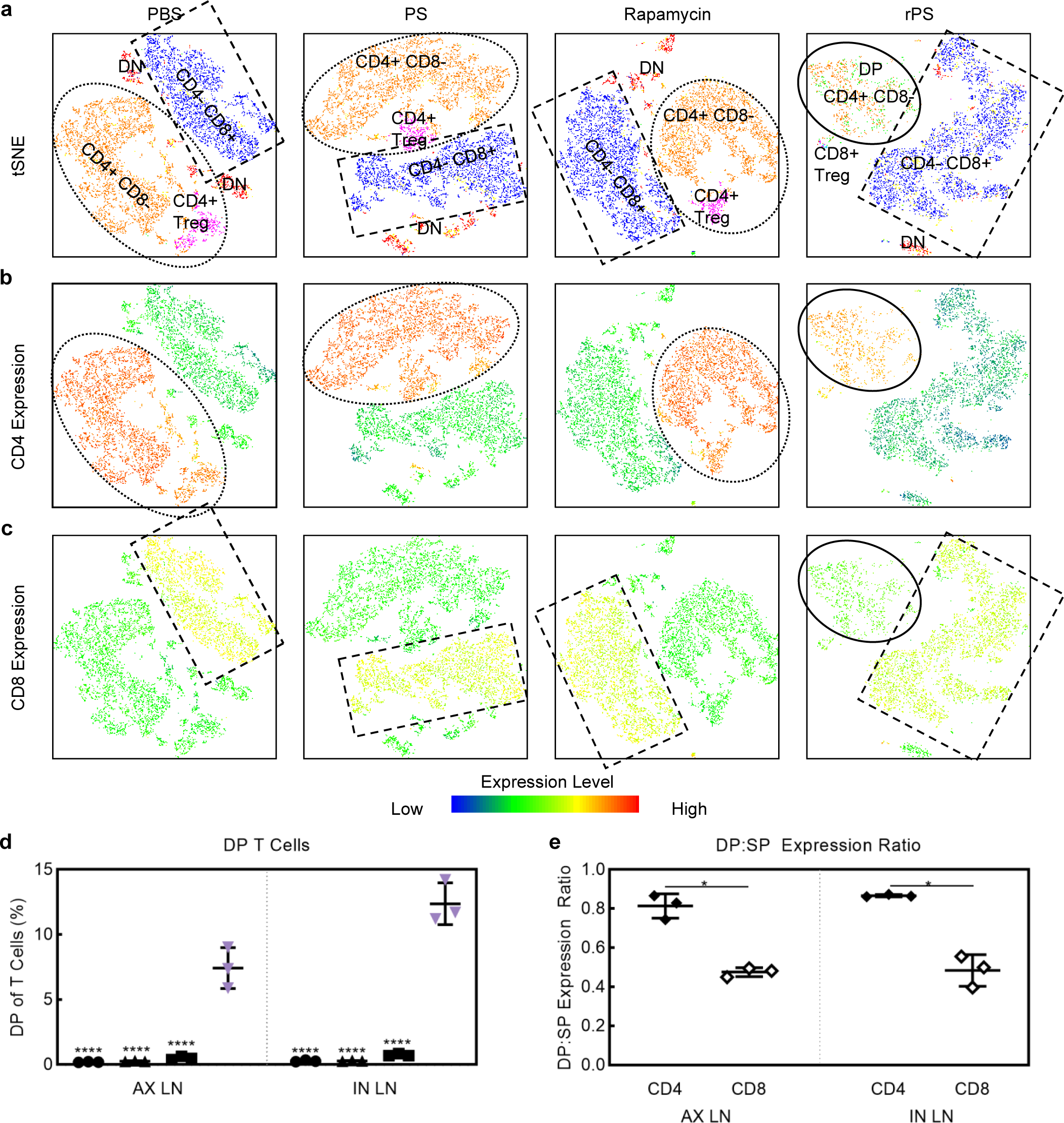
rPS induce DP CD4^bright^ CD8^dim^ T cells with suppressor functions in lymph nodes. Flow cytometry analysis of CD45+ cell populations from mice subcutaneously injected with 1X phosphate buffered saline (PBS;●), polymersomes (PS; ▴), rapamycin (▪) or rapamycin-loaded polymersomes (rPS; 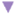) using the standard dosage protocol (11 injections, 1 mg/kg rapamycin or equivalent). **a**, tSNE visualization of CD3+ T cell populations from the axillary lymphocenter (deep axillary/axillary/axial and superficial axillary/brachial lymph nodes; AX LN) with color-coded gated overlays of the previously described cell populations: CD4+ CD8- (orange), CD4-CD8+ (blue), CD4+ CD8+ double positive (DP; green), CD4-CD8-double negative (DN; red), and natural killer (NK; yellow), CD4+ regulatory (CD4+ Treg; magenta) and CD8+ regulatory (CD8+ Treg; light blue). **b,c**, tSNE heatmap statistic of CD4 (**b**) and CD8 (**c**) expression. Dotted line ovals outline CD4+ CD8-T cell populations. Dashed line rectangles outline CD4-CD8+ T cell populations. Solid line ovals outline DP T cell populations. tSNE visualization of CD3+ immune cell populations and CD4 and CD8 heatmap statistics from the subiliac lymphocenter (subiliac/inguinal lymph nodes; IN LN) is shown in **Fig S13. d**, Percentage of DP T cells in AX LN and IN LN. All data are presented as mean percentage (of T cells) ± SD with **** p<0.0001 relative to rPS treatment. Statistical significance was determined by one-way ANOVA with Tukey’s multiple comparisons test. **e**, Ratio of CD4 or CD8 expression by DP:SP (double positive CD4+ CD8+ to single positive CD4+ CD8- or CD4-CD8+) from the AX and IN LN for rPS treated mice. All data are presented as mean ratio ± SD with *p<0.05 relative to rPS treatment. Statistical significance was determined by paired two-tailed t-test. (n = 3 mice/group).

### rPS maintain normoglycemia after fully-MHC mismatched intraportal islet transplantation

*In vivo* assessment of rapamycin redistribution via rPS was conducted using a clinically relevant intraportal (liver) fully-MHC mismatched allogeneic islet transplantation model. Diabetes was induced in C57BL/6J mice via streptozotocin injection. To ensure the most stringent and severe model of T1D, diabetes was defined by blood glucose over 400 mg/dl^23^. A standard dosage protocol known to allow for fully-MHC mismatched allogeneic islet graft viability for more than 100 days was compared to a low dosage protocol (**Fig. 8a**). The standard dosage protocol consisted of 11 injections given daily. The low-dosage protocol consisted of 6 doses given every 3 days (**Fig. 8a**). All doses were equivalent (1 mg rapamycin per kg body weight) (**Fig. 8a**). Diabetic C57BL/6J mice received approximately 200 islets from fully MHC mismatched Balb/c mice in the liver via the portal vein (175 IEQ). Efficacy of the dosing regimen was confirmed by the restoration and maintenance of normoglycemia, confirming survival of the islet graft. As expected, mice that did not receive treatment all experienced graft rejection within 10 days of transplantation (**Fig. 8c,d**) and 71% of mice treated with the standard rapamycin protocol remained normoglycemic 100 days post transplantation (**Fig. 8c,d**). When the low-dosage protocol was used, only 33% of the mice treated with rapamycin remained normoglycemic 100 days post-transplantation, whereas 83% of (all, but one) mice treated with low-dosage rPS had normal blood glucose concentrations (**Fig. 8c,d**). Furthermore, intraperitoneal glucose tolerance test (**IPGTT**), conducted at 30 days post-transplantation showed no difference in islet responsiveness with low dosage rPS treatment as compared to standard dosage rapamycin (**Fig. S22**).

**Fig. 8.**
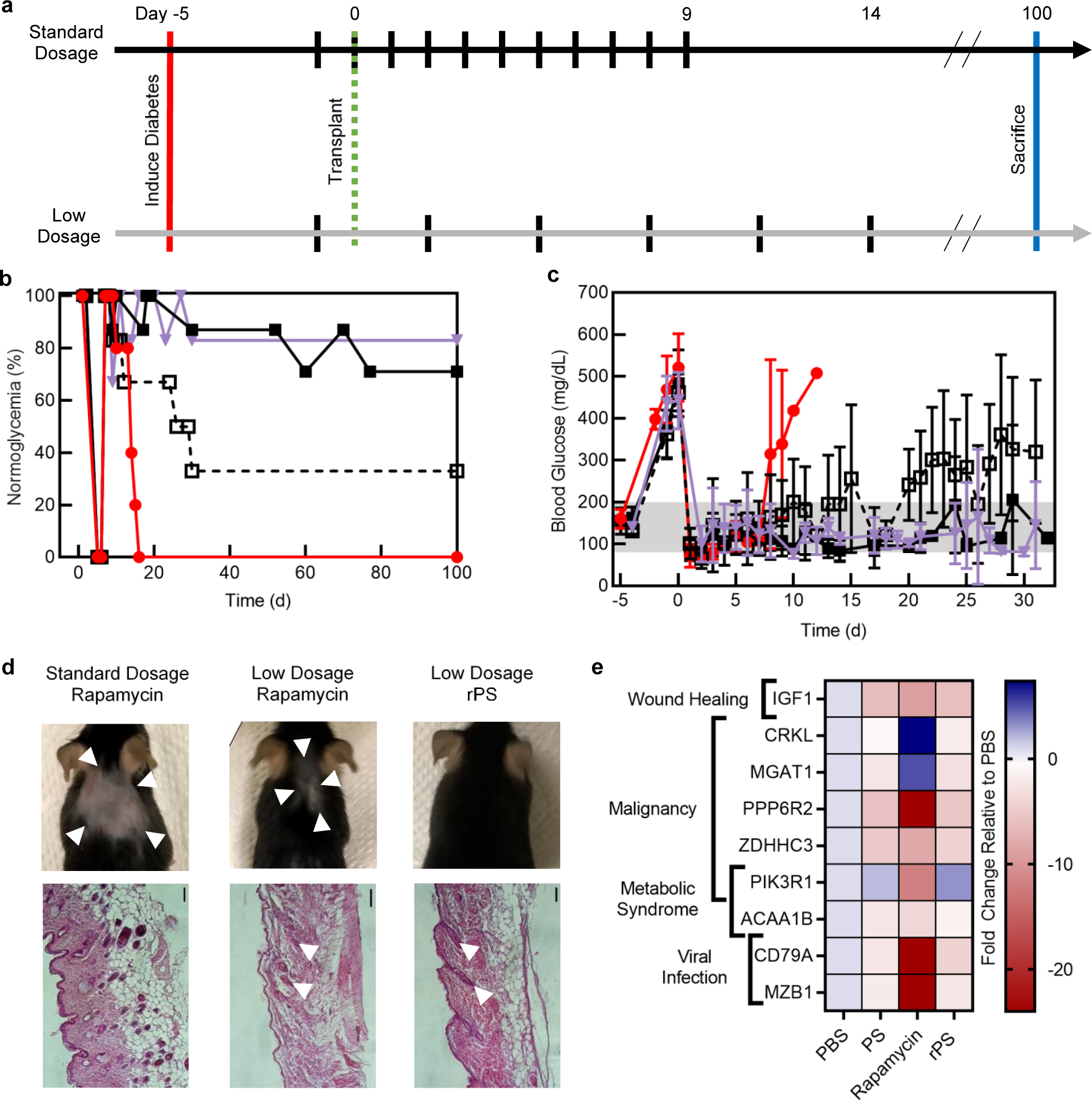
rPS reduce effective drug dose to achieve normoglycemia and mitigate side effects *in vivo*. **a**, Standard dosage and low dosage schemes for rapamycin during allogeneic islet transplantation (day 0) experiment. Diabetes is induced (day -5) via streptozotocin (STZ) injection. The standard dosage protocol consists of 11 injections, given daily starting at day -1. The low dosage protocol consists of 6 injections, given every 3 days, starting at day -1. **b**, Post-transplantation normoglycemia (%) (blood glucose < 200 mg/dl). No treatment 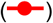; Standard dosage rapamycin 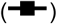; Low dosage rapamycin 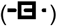; Low dosage rapamycin-loaded polymersomes (rPS) 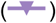. (n=5-7 mice/group). **c**, Post-transplantation blood glucose concentration (mg/dl). No treatment 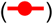; Standard dosage rapamycin 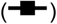; Low dosage rapaymcin 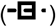; Low dosage rapamycin-loaded polymersomes (rPS;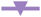). (n=5-7 mice/group). All data are presented as mean glucose concentration ± SD. **d**, Top: Digital photos of subcutaneous injection site on mouse dorsal showing alopecia 30 days after allogeneic islet transplantation by treatment group. Bottom: Hematoxylin and eosin histology of skin taking from mice 100 days post-transplantation with. White arrows show mature hair follicles. Scale bars represent 100 µm. (n 5-7 mice). **e**, Single cell RNA sequencing analysis of macrophages and Tregs from the liver and spleen of genes associated with known side effects of rapamycin, including impaired wound healing, malignancy, metabolic syndrome, and enhanced predisposition to viral infection. (n=3 mice/group). All data are presented as mean fold change.

### Subcutaneous delivery of rapamycin to APCs via rPS avoids rapamycin side effects

We observed that mice treated with free rapamycin experienced injection site alopecia (**Fig. 8d**). Alopecia is a known side effect of rapamycin, impacting approximately 10% of patients^24^. While alopecia was reduced in the low dosage free rapamycin group (**Fig. 8d, S16**), no alopecia was observed in the low dosage rPS group (**Fig. 8e, S17**). Histological analysis confirms our gross observations (**Fig. 8d**). Only immature follicles were identified in the standard rapamycin group (**Fig. 8d**) with some mature follicles present in the low dosage free rapamycin group (**Fig. 8d**). Organized mature follicles were identified in the low dosage rPS group (**Fig. 8d**). Furthermore, single cell RNA sequencing analysis of macrophages and CD4+ Tregs from the spleen and liver demonstrated that rPS mitigated expression of genes associated with rapamycin adverse effects. Specifically, rPS caused less inhibition of insulin-like growth factor 1 (**IGF1**), which is associated with impaired wound healing (**Fig. 8e, Table S1-2**). Oncogenes CRKL (V-Crk Avian Aarcoma Virus CT10) was downregulated with rPS treatment, whereas it was upregulated with rapamycin treatment^25^ (**Fig. 8e, Table S1-2**). Tumor suppressor genes known to be downregulated by rapamycin, including MGAT1 (Mannosyl Glycoprotein Acetylglucosaminyl-Transferase 1)^26^, PIK3R1 (Phosphoinositide-3-Kinase Regulatory Subunit 1)^27^, PPP6R2 (Protein Phosphatase 6 Regulatory Subunit 2)^28^, and ZDHHC3 (Zinc Finger DHHC-Type Palmitoyltransferase 3)^29^ were less inhibited with rPS (**Fig. 8e, Table S1-2**). Furthermore, inhibition of genes associated with the regulation of metabolic processes caused by rapamycin, including ACAA1 (Acetyl-CoA Acyltransferase 1)^30^ and PIK3R1^31^, were reduced when rapamycin is given in rPS form (**Fig. 8e, Table S1-2**). Rapamycin causes downregulation of genes associated with the protective response to viral infection, including CD79A^32^ and MZB1 (Marginal Zone B And B1 Cell Specific Protein)^33^ (**Fig. 8e, Table S1-2**). The inhibition of these viral response genes was not seen with rPS treatment (**Fig. 8e, Table S1-2**).

## Discussion

A grand challenge of pharmaceutical development is to harness the rational engineering of nanoscale drug carriers (i.e. nanocarriers) to selectively modify target cells while minimizing uptake by cells and organs responsible for side effects^34^. By controlling delivery kinetics and target specificity, nanocarriers can alter the interconnected network of cells contributing to observed therapeutic effects, thus significantly changing the therapeutic window and reducing both the dosage and adverse events of a drug during treatment^34^. Effects of changing the network of targeted cells is particularly evident during immunotherapy, where small subsets of immune cells can elicit potent cytokine and T cell responses that can propagate into unique systemic responses. With these concepts in mind, we sought to investigate how subcutaneous delivery and nanocarrier-directed changes in the cellular biodistribution of rapamycin, a common therapeutic that elicits diverse cell-specific effects, could repurpose its mechanism of action at the cellular level to decrease side effects and enhance efficacy.

The targeted cell population and amount of delivered drug are critical considerations for targeted therapies. Rapamycin achieves immunosuppression by directly acting on T cells^3^. However, when given clinically via standard oral administration, the resulting broad biodistribution of rapamycin influences numerous off-target cells and reduces the dose that reaches T cells for desired effects^3,5,35^. Lack of specificity cannot be overcome with increased dosage given that rapamycin is associated with dose-dependent toxicity^5,35^. We have previously shown that giving drugs, including rapamycin, via PS, allows for selective uptake by APCs while avoiding T cells^12,17^. We hypothesized that switching the target cell population from T cells to APCs would change the immunosuppressive mechanism of rapamycin to one that required less dosage and reduced side effects.

Route of administration is another tool that is employed to impact biodistribution and overcome drug-specific barriers to delivery. For example, subcutaneous injection could avoid diet-dependent bioavailability and variable metabolism via CYP3A4 and P-glycoprotein that are associated with orally administered rapamycin^3^. Using these tools—cellular targeting and route of administration—the temporal and both organ and cellular biodistribution of a drug can be precisely manipulated for the desired effect. Herein, we show that subcutaneous delivery of rapamycin via PS creates a rapamycin biodistribution that perturbs the network of inflammatory cells in a manner that supports the survival of transplanted allogeneic islets. While others have attempted to use nanocarriers for altered delivery of rapamycin^35^, to the best of our knowledge, we showcase the first attempt to deliver rapamycin via nanocarriers subcutaneously.

Although drug-loaded PS primarily target APCs within lymphoid organs, as we have previously shown^17^, the effects of subcutaneous administrated rPS span beyond APCs to modulate a diverse network of immune cells. The most profound cellular effects of rPS were seen in the draining AX LN which included an upregulation of APCs and a down regulation of T cells. These rPS-induced modulations of immune cells provide a foundation for an inflammatory environment that is amenable to allogenic islet transplantation. Importantly, we show a downregulation of T cells in immunomodulatory organs and at the site of intraportal islet transplantation—the liver—a key objective of immunosuppressive rapamycin therapy^3^. This was achieved without directly targeting T cells, and instead via enhanced targeting of APCs that dictate T cell function during inflammatory responses^22^. Thus, redistribution of our cellular network via rPS treatment establishes a foundation for cellular immunomodulation.

While direct donor antigen recognition by both CD4+ or CD8+ T cells and indirect presentation of donor antigen to CD8+ T cells contribute to a rejection response, only indirect donor antigen presentation to CD4+ T cells is required for rejection^36^. In this important regard, rPS cause deletion and/or anergy in CD4+ T cells as indicated by the significant reduction in the CD4+ T cell population and reduction in expression of CD3 and CD4^22,37^. The unique combination of costimulation blockade as evidenced by reduced CD40/80/86 expression and enhanced MHC II presentation by APCs is responsible for CD4+ T cell demise.

In the lymph nodes, rPS induces phenotypic changes in the DC population which are amenable to islet transplantation tolerance. Phenotypic changes are enhanced by symbiotic relationships between these DCs and CD8+ T cells to promote a quiescent environment. Along with the overall significant increase in DCs, a significant increase in novel DP CD8+ CD11b+ cDCs was observed. CD11b+ cDCs cross-present antigens to CD4+ T cells and CD8+ cDCs cross-present antigens to CD8+ T cells to induce tolerogenic behavior^9^. The presence of DP cDCs suggests that these cells may have the ability to cross-present donor antigens to both CD4+ and CD8+ T cells or DP CD4+ CD8+ T cells, which are also significantly upregulated in the lymph nodes. Furthermore, tolerogenic tDCs can cause CD8+ T cells to become CD8+ CD25+ FoxP3+ Tregs. CD8+ Tregs have enhanced suppressor capabilities relative to their CD4+ counterparts^38^. The tolerogenic properties of CD8+ Tregs have been shown to prevent graft-versus-host disease and autoimmune diseases^38^. Despite their tolerized state, CD8+ Tregs cells confer immunoprotection against pathogens^38^. In addition, rPS causes a significant downregulation of pDCs, which are known to secrete interferon-gamma and activate cytotoxic CD8+ T cells^39^. Both interferon-gamma secretion and cytotoxic CD8+ T cells are known to damage islet grafts, thus pDC-mediated reduction boosts potential for graft survival^40^. In addition, a significant increase in CD8+ NK T cells was observed. CD8+ NK T cells cause apoptosis of antigen-bearing DCs^41^, thus promoting graft survival. CD8+ NK cells are also known to impede viral infection^42^, a common side effect of immunosuppressive therapy. Checks and balances between DCs and CD8+ T cells prevent cytotoxic activity against the graft.

In addition to DCs, rPS treatment induces suppressor phenotypes in M/Ms. Suppressor M/Ms are notable in blood lymph nodes and spleen. While other treatments confer a M/M population that is dividing between activator and suppressor phenotypes, rPS treatment promotes a single phenotype characterized by its MHC II+ Ly-6C^Lo^ status. Mature MHC II+ Ly-6C^Lo^ M/Ms are a type of patrolling cell that is able to penetrate tissue during steady state conditions. Ly-6C^Lo^ monocytes have the ability to phagocytose both nanoparticles and apoptotic debris^19^. This non-classical monocyte population has a dual-fold advantage for transplantation applications, in which it supports an anti-inflammatory phenotype amenable to graft tolerance^43^ and it has been shown to aid in the prevention of viral infections^44^. These monocytes have the ability to cross present the apoptotic debris to CD8+ T cells and tolerize the CD8+ T cell, suppressing antigen specific responses^19^, in a similar manner to that of CD8+ cDCs^9^.

The tolerogenic effects of rPS-treated APCs span beyond CD4+ and CD8+ T cells to create a hospitable environment of the islet graft. Niche T cell populations also make an important contribution to the congenial environment observed with rPS immunomodulatory therapy. For example, the upregulation of novel DP CD4+ CD8+ T cells in the lymph nodes is observed. Controversy has surrounded DP T cells as both suppressive and cytotoxic functions have been demonstrated^45,46^. This is because while CD4^dim^ CD8^bright^ cells are cytotoxic^45^, CD4^bright^ CD8^dim^ DP T cells are anti-inflammatory^46,47^. tSNE visualization in combination with CD4 and CD8 expression level analysis helped to revel that rPS confer CD4^bright^ CD8^dim^ DP T cells with known suppressor function, such as secreting anti-inflammatory cytokines^47^. Interestingly, these cells also show enhanced responsiveness during infection, for example activating effector cells in the case of human immunodeficiency virus^47^.

Our use of the liver transplantation site is critical for translation of murine studies as the commonly used kidney capsule is not a feasible site for human islet transplantation^23^. Kidney capsule transplantation fails to expose the islets to the immune environment of the liver^23^. For example, islets transplanted to the kidney capsule are not exposed to blood to induce the instant blood-mediate inflammatory reaction (IBMIR)^23^. Additionally, the exposure of the islets to immunosuppressive drugs differs between the liver and kidney capsule transplantation site^23^. When islets are infused into the vasculature of the liver, they first come into contact with neutrophils. Furthermore, rPS treatment significantly reduces the neutrophil population in blood and liver and downregulates expression of CD11b (**Fig. S15**). CD11b is critical for neutrophil migration^48^. Graft infiltrating neutrophils have been shown to cause transplant failure^49^, thus reduction in this cell type and reduced mobility may contribute to enhanced graft survival. With reduced CD11b, neutrophils may show decreased ability to reach islets and infiltrate the graft. Furthermore, MHC molecules are the most significant allo-antigens involved in graft rejection, thus using a fully MHC-mismatched model is critical for rigorous assessment of allogeneic transplantation. Furthermore, it is important to note that all combinations of fully mismatched mouse models confer the same potency and kinetics of allo-immune response. We utilized the combination of Balb/c islets transplanted into C57BL/6J recipient mice, which provides the greatest challenge to islet survival and normoglycemia restoration^23^. Utilizing excess islets can delay the graft rejection, giving a false sense of maintained normoglycemia and immunosuppression. While other models use up to 1000 islet equivalents (**IEQ**)^2,50^, our model uses a minimal islet mass of only ∼200 murine islets (∼175 IEQ).

The translatability of the model is enhanced by employing subcutaneous injection, which supports lymphatic drainage and simplifies therapeutic administration (**Fig. 3a,b**)^10^. Furthermore, this route of administration overcomes some of the challenges that have historically plagued the oral, nanocrystal formulation of rapamycin, Rapamune®^35^. Specifically, subcutaneous delivery of rapamycin improves bioavailability over oral delivery as first pass metabolism and p-glycoprotein efflux are avoided^35^. Variability of bioavailability as food intake is not an influencing factor for subcutaneous injection. Many murine studies involving rapamycin use intraperitoneal injection^2^, however this route is not easily translatable to humans. Finally, subcutaneous injection is advantageous over intravenous infusion as patients can perform the injection in their own home, as opposed to requiring the services of a health care professional. In summary, this work lays a foundation for a novel method of repurposing clinically relevant drugs by targeting them to selected cell populations to rationally modulate the therapeutic mechanism of action for enhanced efficacy while mitigating adverse effects.

## Materials and Methods

### Animals

8 to 12-week-old, male C57BL/6J and Balb/c mice were purchased from Jackson Labs. Mice were housed in the Center for Comparative Medicine at Northwestern University. All animal protocols were approved by Northwestern University’s Institutional Animal Care and Use Committee (**IACUC**).

### Materials

Unless explicitly stated below, all reagents and chemicals were purchased from Sigma-Aldrich.

### Polymer Synthesis

PEG-*b*-PPS was synthesized as previously described by us^12^. In brief, methyl ether PEG (MW 750) was functionalized with mesylate. The mesylate was reacted with thioacetic acid to form PEG-thioacetate and then base activating the thioacetate to form a thiolate anion and initiate ring opening polymerization of propylene sulfide. Benzyl bromide was used as an end-capping agent to form PEG_17_-*b*-PPS_30_-Bz or the thiolate anion was protonated to form PEG_17_-*b*-PPS_30_-SH. The polymer was characterized by H-NMR and gel permeation chromatography (**GPC**).

### Nanocarrier Formulation

PS were formed via thin film hydration, as previously described^12,17^. In brief, 20 mg of PEG_17_-*b*-PPS_30_-Bz was weighted in a sterilized 1.8 ml glass HPLC vial. 750 ul of dichloromethane (**DCM**) was added to the vial. To form, rPS 0.5 mg of rapamycin, dissolved at 25 mg/ml in ethanol, was also added. The vial was desiccated to remove the DCM. Next, 1 ml of PBS was added to the vial. The vials were shaken at 1500 rpm overnight. PS were extruded multiple times first via 0.2 um and then 0.1 um syringe filters. Excess rapamycin was removed via size exclusion chromatography using a Sephadex LH-20 column with PBS.

### Nanocarrier Characterization

#### DLS

DLS measurements were performed on a Nano 300 ZS Zetasizer (Malvern) and were used to determine nanocarrier diameter distribution and corresponding polydispersity index.

#### cryoTEM

200-mesh lacey carbon grids were glow-discharged for 30 seconds in a Pelco easiGlow glow-discarger at 15mA with a chamber pressure of 0.24 mBar. 4 µL of sample was then pipetted onto the grid and plunge-frozen into liquid ethane in a FEI Vitrobot Mark III cryo plunge freezing device for 5 seconds with a blot offset of 0.5mm. Grids were then loaded into a Gatan 626.5 cryo transfer holder, imaged at –172 °C in a JEOL JEM1230 LaB6 emission TEM at 100kV, and the data was collected on a Gatan Orius 2k x 2k camera.

#### SAXS

SAXS was performed at Argonne National Laboratory’s Advanced Photo Source with collimated X-rays (10 keV; 1.24 Å). Data reduction was performed using Primus software and modeling was performed using SASView.

### Quantification of Rapamycin Loading^12^

rPS nanocarriers (50 ul) were lyophilized and re-dissolved in HPLC grade dimethylformamide (**DMF**). Salts were removed via centrifugation at 17,000 g for 10 minutes. Rapamycin content of the nanocarriers was characterized via HPLC (Thermo Fisher Dionex UltiMate 3000) using an Agilent Polypore 7.5 ⨯ 300 mm column and an Agilent Polypore 7.5 ⨯ 50 mm guard column. The system was housed at 60°C. DMF (0.5 ml/minute) was used as the mobile phase. Rapamycin was detected at 270 nm. Thermo Scientific Chromeleon software was used for analysis. The concentration of rapamycin was characterized via the area under the curve in comparison to a standard curve of rapamycin concentrations.

### Rapamycin Stability in Nanocarrier

rPS formulations were fabricated as previously described. Formulations were stored at 4°C in glass scintillation vials. At various time points, the formulations were vortexed, 1 ml samples were transferred to Millipore Amicon Ultra Centrifuge 10,000 NMWL Tubes and centrifuged at 4000 g in a swinging bucket rotor to remove unloaded drug. The retentate was brought back up to its original volume using PBS. Quantification of rapamycin was performed as previously described.

### Immunomodulation Study

Healthy C57BL/6J mice were subjected to a “standard dosage regime.” Animals were injected subcutaneously for 11 days with rapamycin (in 0.2% CMC) or rPS at a dose of 1 mg/kg. Equivalent dose of PBS or PS were injected as controls. After 11 days, the mice were sacrificed. Blood, liver, AX LN, IN LN and spleen were collected and processed for flow cytometry.

### Flow cytometry

Blood was spun down at 3000 g for 25 minutes to separate the plasma and blood cells. The blood cells were treated with 1X red blood cell lysis buffer (Fisher) for 5 minutes on ice, washed with PBS, and spun down, thrice. The liver was minced, treated with collagenase for 45 minutes at 37 °C, processed through a 70 nm filter, and then treated with 1X red blood cell lysis buffer (Fisher) for 5 minutes on ice, washed with PBS and spun down.

The spleen was processed through a 70 nm filter and treated with 1X red blood cell lysis buffer (Fisher) for 5 minutes on ice, washed with PBS and spun down. Lymph nodes were passed through a 70 nm filter, washed with PBS and spun down. All cells were resuspended in a cocktail of Zombie Near Infrared (BioLegend) for viability and anti-mouse CD16/CD32 for FcR blocking with BD Brilliant Violet cell staining buffer and incubated at 4 °C for 15 minutes. Next, an antibody cocktail consisting of Pacific Blue anti-mouse CD11c (BioLegend), BV480 anti-mouse NK1.1 (BD), BV510 anti-mouse CD19 (BioLegend), BV570 anti-mouse CD3 (BioLegend), BV650 anti-mouse F4/80 (BioLegend), BV650 anti-mouse MHC II (IA-IE) (BioLegend), BV711 anti-mouse Ly-6C (BioLegend), BV750 anti-mouse CD45R/B220 (BioLegend), BV785 anti-mouse CD11b (BioLegend), AF532 anti-mouse CD8a (Invitrogen), PerCP-Cy5.5 anti-mouse CD45 (BioLegend), PerCp-eFluor711 anti-mouse CD80 (Invitrogen), PE-Dazzle594 anti-mouse CD25 (BioLegend), PE-Cy5 anti-mouse CD4 (BioLegend), PE-Cy7 anti-mouse CD169 (BioLegend), APC anti-mouse FoxP3 (Invitrogen), AF647 anti-mouse CD40 (BioLegend), APC-R700 anti-mouse Ly-6G (BioLegend), and APC/Fire 750 anti-mouse CD86 (BioLegend) was added to the cells and incubated for 20 minutes at 4 °C. The cells were washed with PBS, fixed and permeabilized using a FoxP3 Fix/Perm Kit (BioLegend), according to the manufacturer’s protocol. Next, anti-mouse FoxP3 was added and incubated for 30 minutes in the dark at room temperature. Finally, cells were washed twice with PBS and resuspended in cell buffer. The cells were analyzed on an Aurora flow cytometer (CyTek). Spectral unmixing was performed using SpectroFlo (CyTek) and analysis was performed using FloJo software. Gating was performed as outlined in **Fig. S4**^51,52^.

### tSNE

For each analysis, FlowJo’s DownSample plugin was used to randomly select an equal number of events from each cell population (CD45+, CD45+ CD3+, CD45+ CD19+, CD45+ CD11b+, or CD45+ CD11c+) of every sample. The purpose of DownSample was to both normalize the contribution of each mouse replicate and reduce computational burden. Next, samples from mice that underwent the same treatment and same cell population were concatenated. The tSNE plugin was run on concatenated samples using the Auto opt-SNE learning configuration with 3000 iterations, a perplexity of 50 and a learning rate equivalent to 7% of the number of events^53^. The KNN algorithm was set to exact (vantage point tree) and the Barnes-Hut gradient algorithm was employed.

### ICG Biodistribution

ICG PS were formed using thin film rehydration, as previously described^17^. In brief, 20 mg of PEG_17_-*b*-PPS_30_-Bz was weighted in a sterilized 1.8 ml glass HPLC vial. 750 ul of DCM was added to the vial. The vial was desiccated to remove the DCM. Next, 1 ml of 0.258 mM ICG in PBS was added to the vial. The vials were shaken at 1500 rpm overnight. PS were extruded multiple times first via 0.2 um and then 0.1 um syringe filters. Float-A Lyzer G2 Dialysis devices (Fisher) were used to remove unloaded ICG. ICG loading was quantified relative to standards composed of known amounts of polymer and ICG in a 1:33 molar ratio using absorbance at 820 nm as previously described by our group^17^. C57BL/6J mice received subcutaneous injections of either free ICG (in PBS) or ICG-PS. ICG concentration was matched at 50 ug/ml. The injection volume was 150 ul. At 2, 24- and 48-h post-injection, the mice were sacrificed, blood was collected via cardiac puncture, and perfusion was performed using heparinized PBS. Liver, spleen, kidneys, heart and lung were harvested and imaged via IVIS Lumina with an excitation wavelength of 745 nm, an emission wavelength of 810 nm, an exposure time of 2 seconds and a f/stop of 2.

### Rapamycin Biodistribution

Mice were injected with rapamycin (in 0.2% CMC) or rPS at 1 mg/ml and sacrificed at the following time points: 0.5, 2, 8, 16, 24, and 48 h. Urine was collected via metabolic cages during the duration between injection and sacrifice for the 8, 16, 24 and 48-h timepoints. The following tissues and/or organs were collected: blood, brain, fat pad, heart, kidneys, liver, lungs, AX LN, IN LN, and spleen. Rapamycin was extracted from blood and urine using a solution of methanol and acetonitrile (50:50 v/v) doped with rapamycin-D3 (Cambridge Isotope Laboratories) as an internal standard. Tissue samples were homogenized in homogenization tubes prefilled with stainless steel ball bearings (Sigma) using a solution of phosphoric acid (8%), acetonitrile and acetic acid (30:67.2:2.8 v/v/v). After homogenization, tissue samples were also doped with rapamycin-D3. All samples were precipitated via incubation at -20 °C, followed by centrifugation. The supernatant was collected and LC-MS/MS (Shimadzu LC-30AD pumps; SIL-30ACMP autosampler; CBM-20A oven; Sciex Qtrap 6500) was used to determine rapamycin concentration. Rapamycin had a retention time of 2.7 minutes. Rapamycin-D3 had a retention time of 3.0 minutes.

### Allogeneic Islet Transplantation

Diabetes was induced via streptozotocin (**IP**; 190 mg/kg) injection five days prior to transplantation and confirmed via hyperglycemia (blood glucose > 400 mg/dl). Starting the day prior to transplantation, mice were injected with PBS, PS, rapamycin, or rPS at 1 mg/kg (or equivalent) in accordance with a standard dosage (11 doses, given daily) or a low dosage (6 doses, given every 3^rd^ day). On the day of transplantation, islets were isolated from Balb/c mice via common bile duct cannulation and pancreas distension with collagenase. Islets isolated from two donors (∼200 mouse islets, ∼175 IEQ) were transplanted to C57B6/J recipients via the portal vein. Body weight and blood glucose concentration were monitored closely for 100 days post-transplantation. Intraperitoneal glucose tolerance test (**IPGTT**) was performed one-month post transplantation. The animals were fasted for 16 h before being injected intraperitoneally with 2 g dextrose (200 g/L; Gibco) per kg body weight. Blood glucose concentrations were measured at 0, 15, 30, 60- and 120-minutes post-injection.

### Alopecia Assessment

Dorsal photos were taken weekly to assess for alopecia. At 100-days post-transplantation, the mice were euthanized, and skin samples were excised in the dorsal region at the subcutaneous injection site. Skin samples were placed in cassettes, fixed in 4% paraformaldehyde, and embedded in paraffin. Tissue blocks were sectioned at a thickness of 5 nm and stained with hematoxylin and eosin (**H&E**). Digital images were taken on a Nikon microscope.

### Single Cell RNA Sequencing

Healthy C57BL/6J mice were subjected to a “standard dosage regime.” Animals were injected subcutaneously for 11 days with rapamycin (in 0.2% CMC) or rPS at a dose of 1 mg/kg. Equivalent dose of PBS or PS were injected as controls. After 11 days, the mice were sacrificed, and the liver and spleen were excised. The organs were processed as was done for flow cytometry. CD4+ Tregs and macrophages were isolated using magnetic sorting (MojoSort; BioLegend). Briefly, cells were first incubated in a cocktail of PE anti-mouse CD169 and PE anti-mouse F4/80 antibodies (BioLegend). After washing, incubation in anti-PE nanobeads (BioLegend) occurred. Macrophages were magnetically sorted from non-macrophages. The non-macrophages cell fraction was then incubated in mouse CD4+ T cell isolation biotin-antiboy cocktail (BioLegend) and sorted. The CD4+ T cell fraction was then incubated in APC anti-mouse CD25 antibody (BioLegend), followed by washing, incubation in anti-APC nanobeads (BioLegend) and sorting. RNA was isolated from separated macrophages and CD4+ Tregs using RNeasy Mini Kit with DNAse digestion (Qiagen). Samples were frozen and shipped to Admera Health where they underwent library preparation using the Lexogen 3’ mRNA-Seq Library Prep Kit FWD HT (Lexogen) and were sequenced on an Illumina sequencer (HiSeq 2500 2 ⨯ 150 bp). For each pair, Read 2 was discarded and only Read 1 was used for downstream data analysis. Sequencing quality was analyzed with FastQC v0.11.5^54^ and reads were trimmed and filtered with Trimmomatic v0.39^55^. One sample from the spleen T cell PBS treatment group and one sample from the spleen T cell rapamycin treatment group were discarded due to low sequencing quality. Reads were aligned with STAR v2.6.0a ^56^ to the GRCm38.p6 mouse reference genome primary assembly using the GRCm38.p6 mouse reference primary comprehensive gene annotation (https://www.gencodegenes.org/mouse/). Quantification and differential expression was performed with Cuffdiff from Cufflinks v2.2.1^57-59^ again using the GRCm38.p6 mouse reference primary comprehensive gene annotation and a 0.05 FDR. Detailed settings for each software are included in Table S1. The raw data displayed in **Fig. 8e** broken down by cell type is in **Table S2**.

## Acknowledgements

Alex D. Jerez designed and created the illustration in **Fig. 1**. Modifications were made by J.B. This material is based upon work supported by the National Science Foundation Graduate Research Fellowship under Grant No. DGE-1842165. This work made use of funding from the Center for Advanced Regenerative Engineering (**CARE**), the Flow Cytometry Facility at the University of Chicago, the Integrated Molecular Structure Education and Research Center (**IMSERC**) at Northwestern University, which has received support from the Soft and Hybrid Nanotechnology Experimental (**SHyNE**) Resource (NSF ECCS-1542205), the State of Illinois, and the International Institute for Nanotechnology (**IIN**), the Northwestern University Center for Advanced Molecular Imaging (**CAMI**), which is generously supported by NCI CCSG P30 CA060553 awarded to the Robert H Lurie Comprehensive Cancer Center. the BioCryo facility of Northwestern University’s NU*ANCE* Center, which has received support from the SHyNE Resource (NSF ECCS-1542205); the MRSEC program (NSF DMR-1720139) at the Materials Research Center; the International Institute for Nanotechnology (**IIN**); and the State of Illinois, through the IIN. It also made use of the CryoCluster equipment, which has received support from the MRI program (NSF DMR-1229693).

## Author Contributions

J.B. designed the experiments with the assistance of S.D.. J.B., X.Z., S.B., M.F., C.B., and H.H. performed the experiments. R.R. performed computational analysis on the RNA sequencing data with oversight from LA. JB analyzed the data and composed the manuscript. E.S. and G.A. supervised the study.

## Competing Interests

J.B., S.D., E.S., and G.A. are coinventors on a patient application related to the work presented in this manuscript.

## Notes

### Competing Interest Statement

J.B., S.D., E.S., and G.A. are coinventors on a patent application related to the work presented in this manuscript.

### Summary of Updates

The author list, figures and results were updated.

