## Supplemental data for "Subcutaneous nanotherapy repurposes the immunosuppressive mechanism of rapamycin to enhance allogeneic islet graft viability"

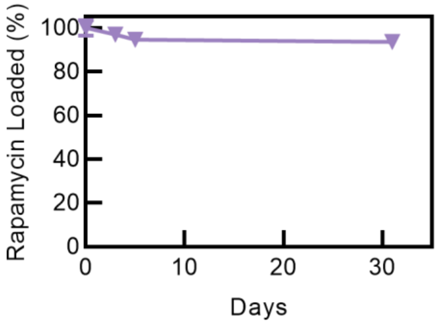

**Fig. S1 | Characterization of rapamycin-loaded polymersomes (rPS) encapsulation stability.** rPS were fabricated, unencapsulated rapamycin was removed and rPS samples were stored at 4 °C in phosphate buffered saline (PBS) at a concentration of 0.125 mg rapamycin/ml. At various time points, released rapamycin was removed and loaded rapamycin concentration was assessed via HPLC-UV. (n = 3-5).

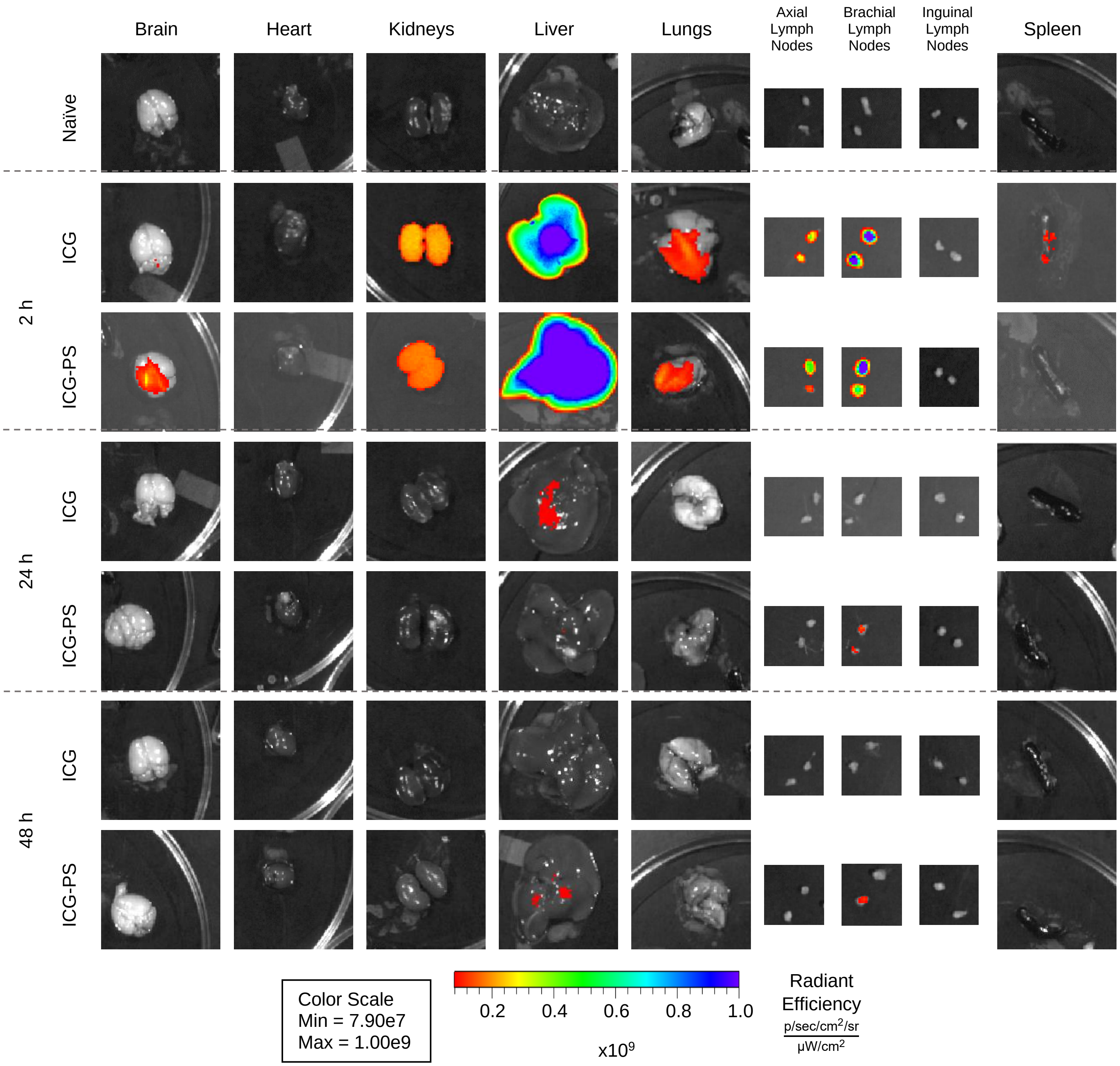

**Fig. S2 | Biodistribution of indocyanine green (ICG) dye and ICG loaded in to polymersomes (ICG-PS).** Mice were subcutaneously injected with ICG-PS or ICG and sacrificed at 2, 24 and 48 hours after injection (n = 5 mice). IVIS was performed on extracted organs to quantify ICG dye.

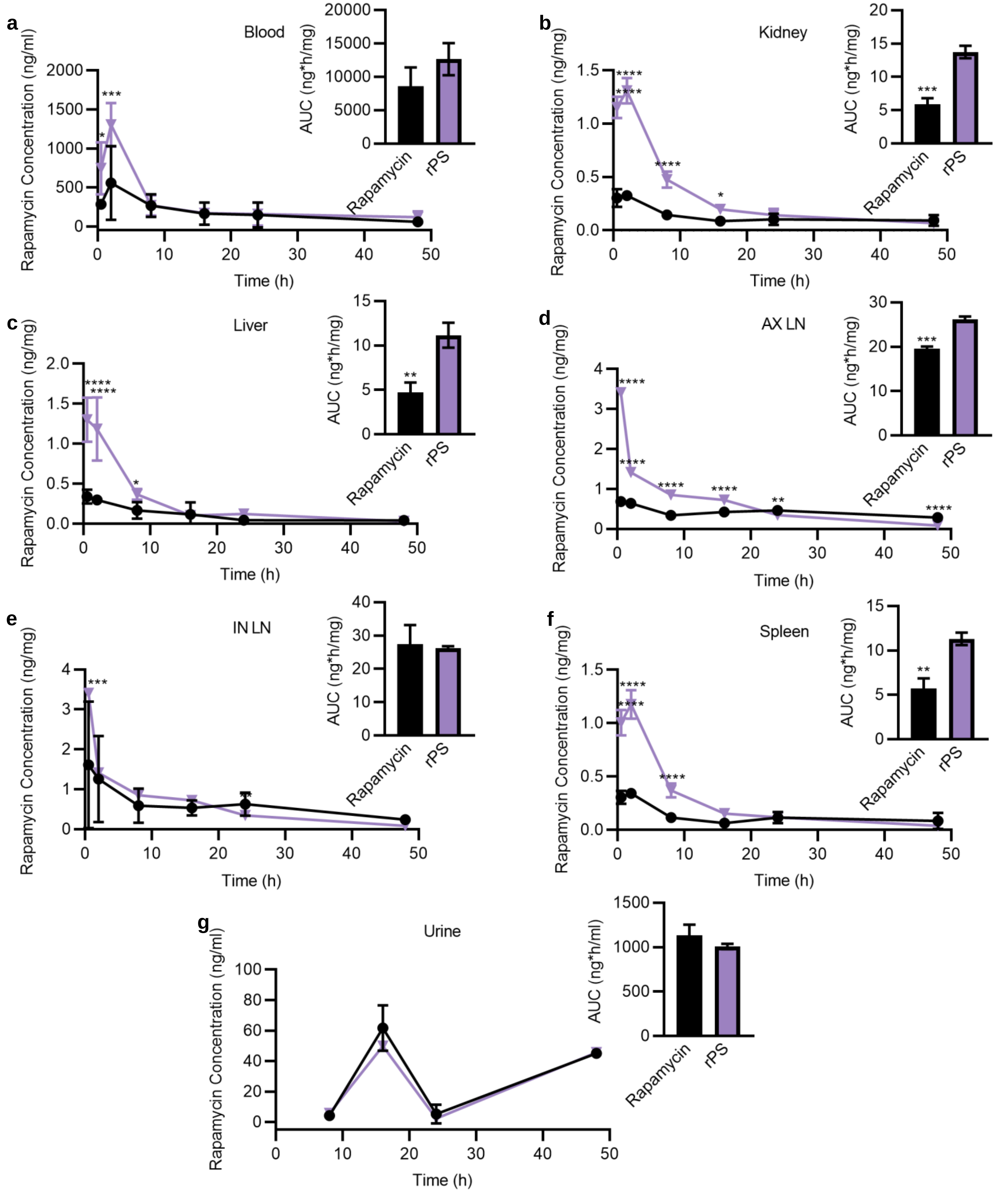

**Fig. S3 | Biodistribution of rapamycin by formulation.** Rapamycin concentration in the **a,** blood, **b,** kidneys, **c,** liver, **d,** axillary lymphocenter (deep axillary/axillary/axial and superficial axillary/brachial lymph nodes; AX LN), **e,** subiliac lymphocenter (subiliac/inguinal lymph nodes; IN LN) **f,** spleen and **g,** urine 0.5 h, 2 h, 8 h, 16 h, 24 h and 48 h after a single subcutaneous injection of rapamycin (in 0.2% carboxymethyl cellulose (CMC)) (—■—) or rPS (—▼—). Rapamycin concentration was also analyzed in the lungs, brain, heart, and fat; concentrations were below 1 ng/mg for both rapamycin and rPS at all timepoints. Analysis was performed using LC-MS/MS. (n = 3 mice/group).

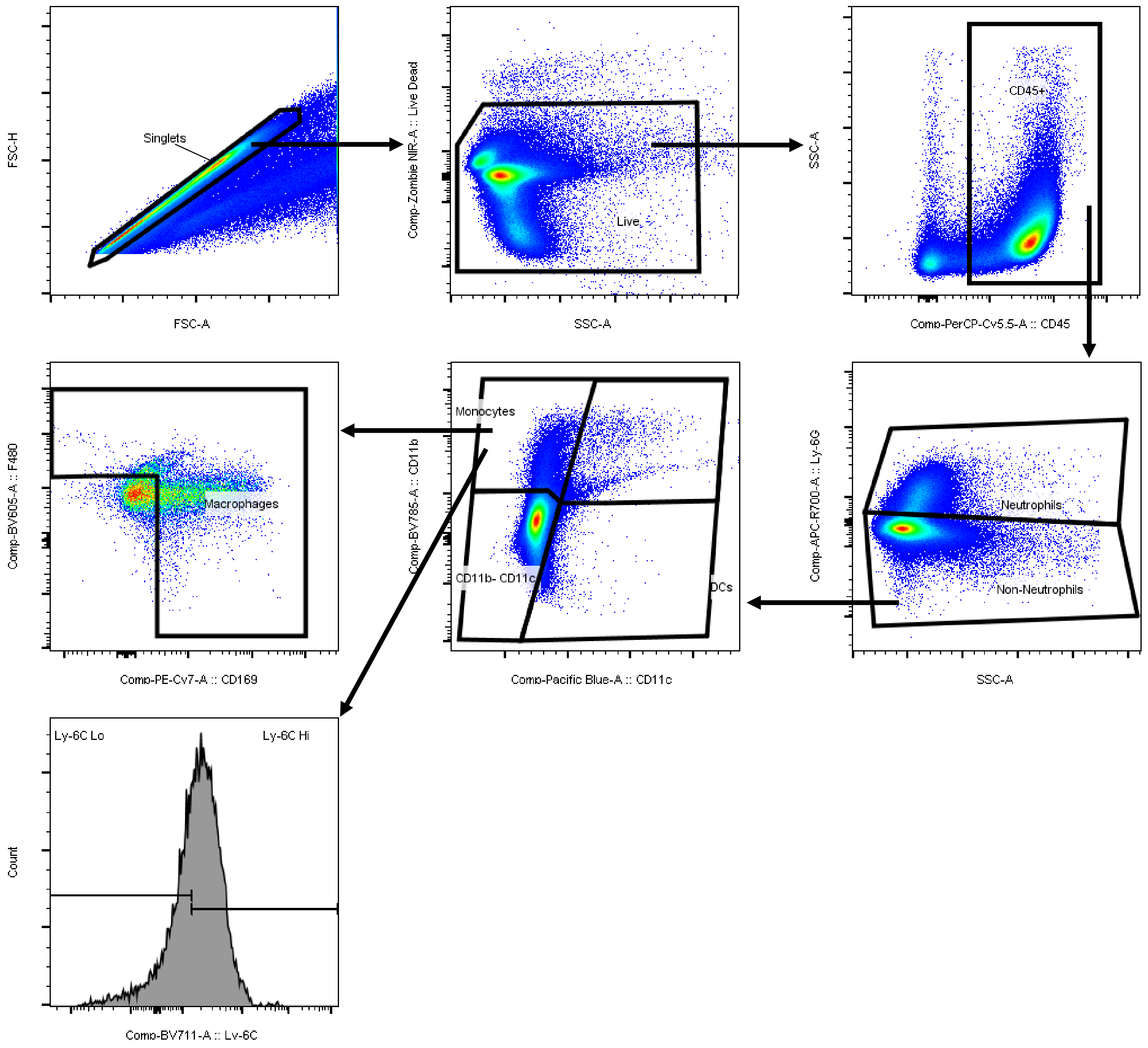

**Fig. S4 | Gating strategy for cell populations in flow cytometry studies.** Representative pseudocolor plots and histograms are displayed from an example mouse lymph node.

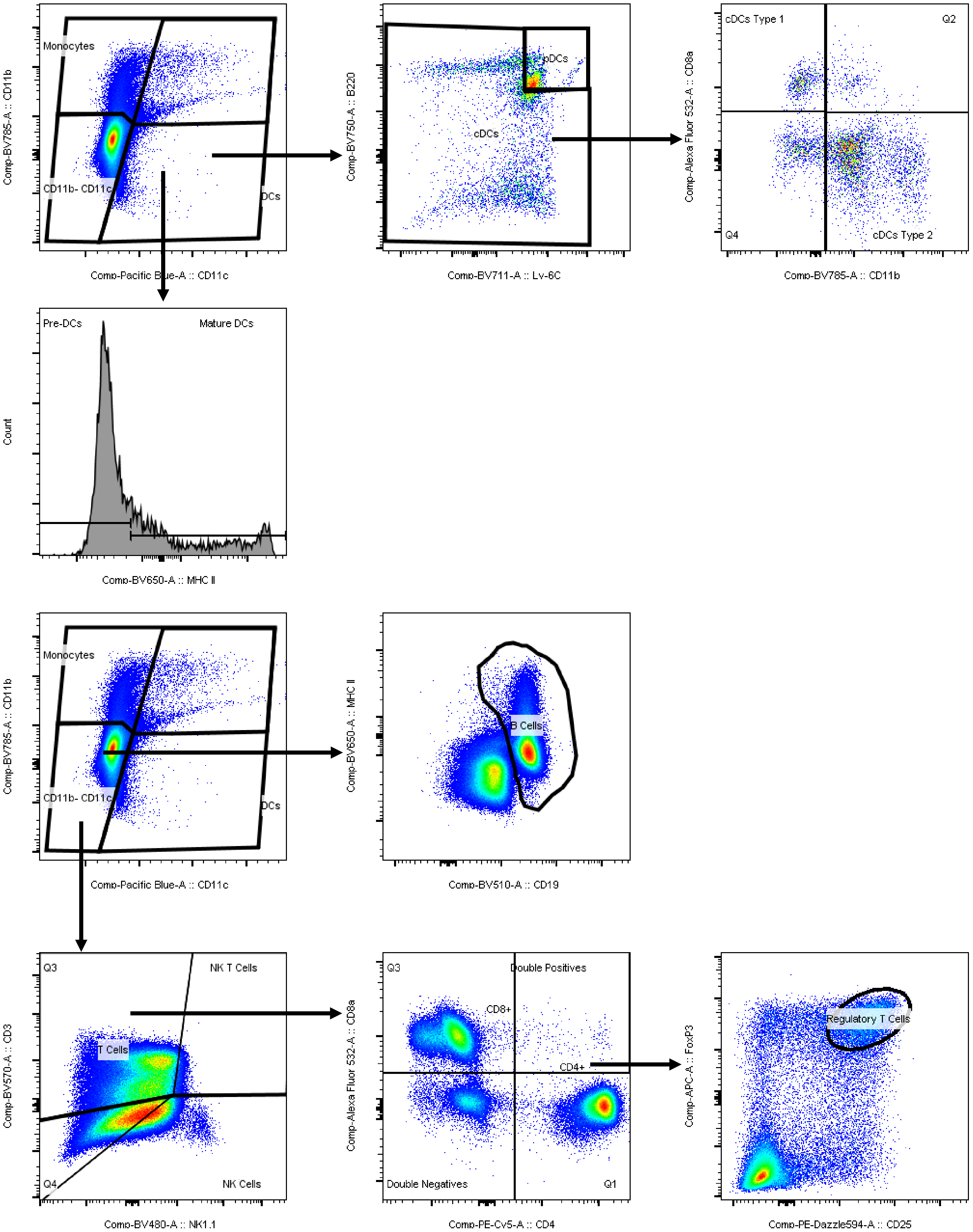

**Fig. S4 (continued) | Gating strategy for cell populations in flow cytometry studies.** Representative pseudocolor plots and histograms are displayed from an example mouse lymph node.

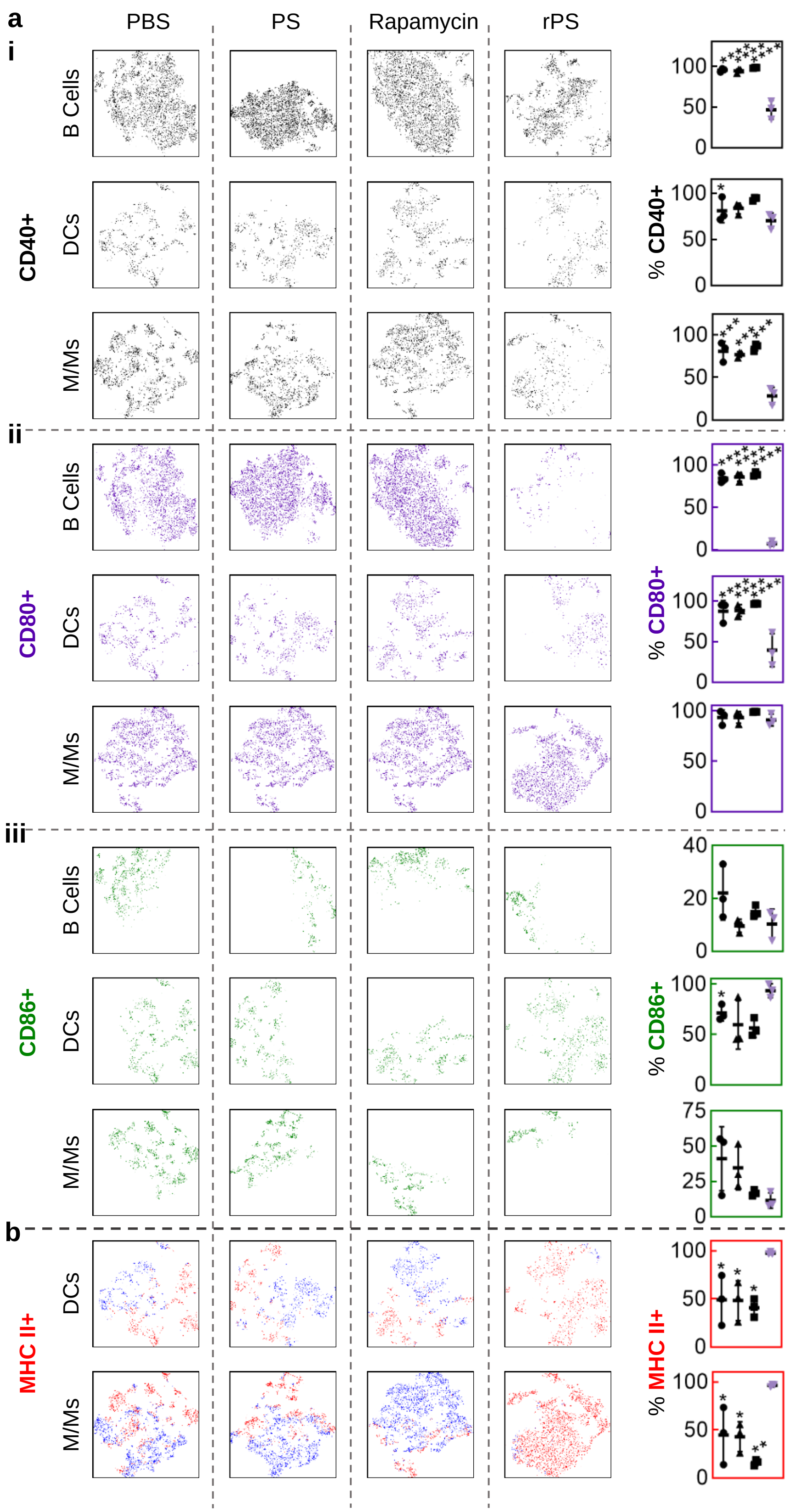

**Fig. S5 | Coreceptor and MHC II expression in blood.** Flow cytometry analysis of CD45+ cell populations from mice subcutaneously injected with phosphate suffered saline (PBS; ●), polymersomes (PS; ▲), rapamycin (■) or rapamycin-loaded polymersomes (rPS; ▼) using the standard dosage protocol (11 injections, 1 mg/kg rapamycin or equivalent). **a**, tSNE visualization of coreceptor CD40 (**a i**; black), CD80 (**a ii**; purple) and CD86 (**a iii**; forest green) positive B cells, dendritic cells (DCs) and monocyte-and-macrophage-linage cells (M/Ms). **b,** tSNE visualization of major-histocompatibility complex (MHC) II presentation (MHC II+ red; MHC II- blue) on DCs and M/Ms. All data are presented as mean percentage (of DCs or M/Ms) ± SD with *p<0.05; ** p<0.01; *** p<0.001; **** p<0.0001 relative to rPS treatment. Statistical significance was determined by one-way ANOVA with Tukey's multiple comparisons test.

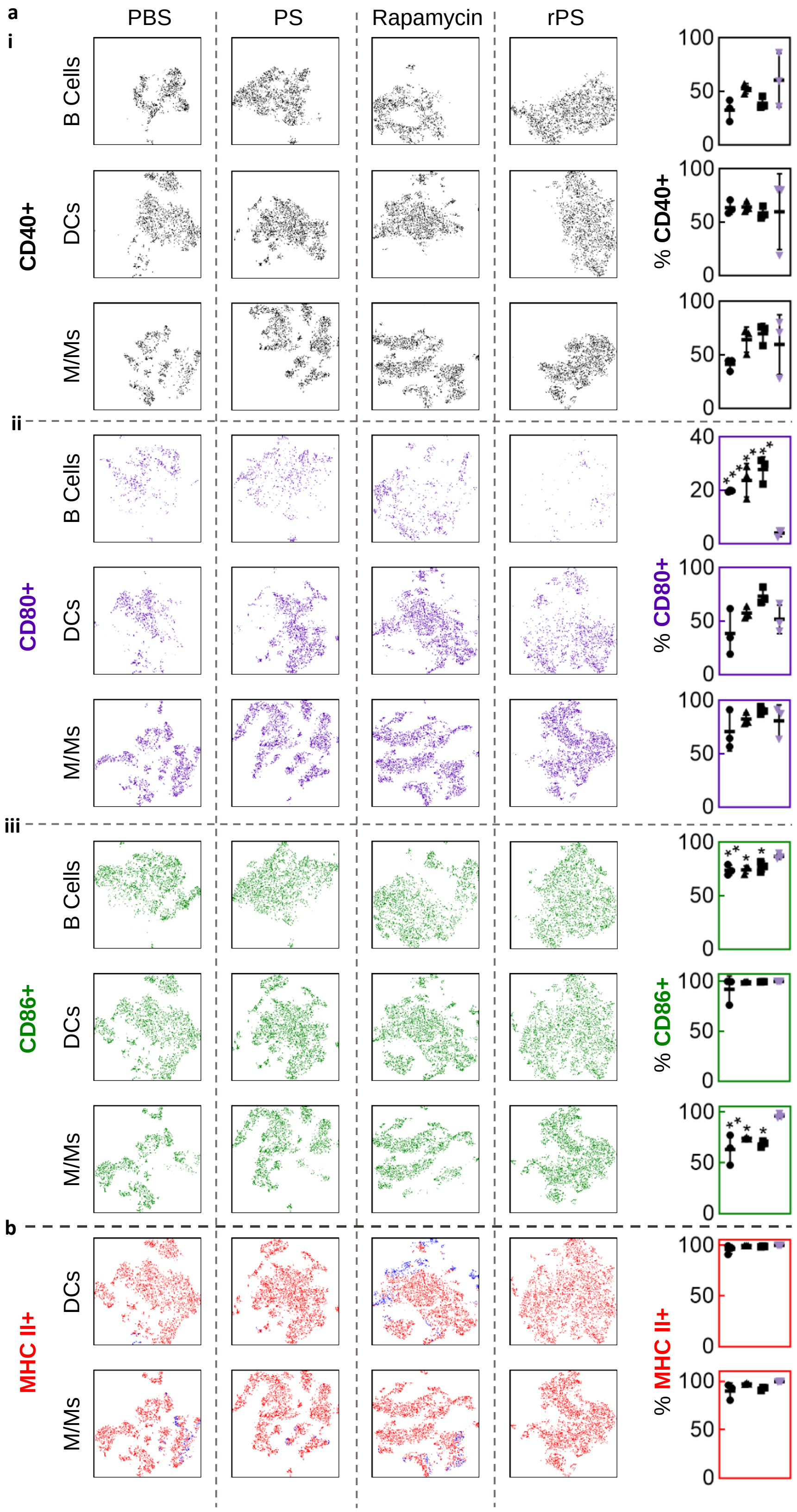

**Fig. S6 | Coreceptor and MHC II expression in liver.** Flow cytometry analysis of CD45+ cell populations from mice subcutaneously injected with phosphate suffered saline (PBS; ●), polymersomes (PS; ▲), rapamycin (■) or rapamycin-loaded polymersomes (rPS; ▼) using the standard dosage protocol (11 injections, 1 mg/kg rapamycin or equivalent). **a**, tSNE visualization of coreceptor CD40 (**a i**; black), CD80 (**a ii**; purple) and CD86 (**a iii**; forest green) positive B cells, dendritic cells (DCs) and monocyte-and-macrophage-linage cells (M/Ms). **b,** tSNE visualization of major-histocompatibility complex (MHC) II presentation (MHC II+ red; MHC II- blue) on DCs and M/Ms. All data are presented as mean percentage (of DCs or M/Ms) ± SD with *p<0.05; ** p<0.01; *** p<0.001; **** p<0.0001 relative to rPS treatment. Statistical significance was determined by one-way ANOVA with Tukey's multiple comparisons test.

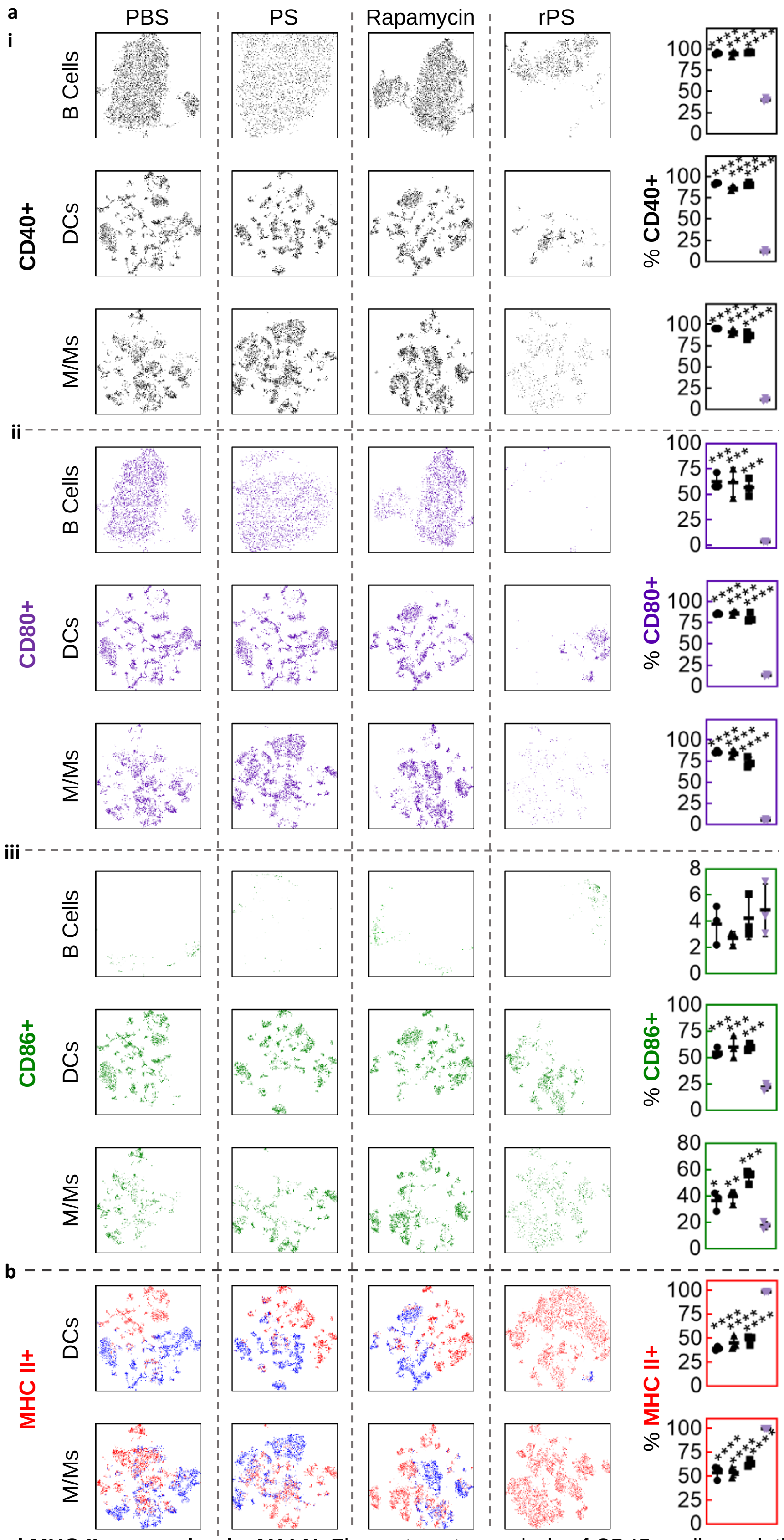

**Fig. S7 | Coreceptor and MHC II expression in AX LN.** Flow cytometry analysis of CD45+ cell populations from mice subcutaneously injected with phosphate suffered saline (PBS; ●), polymersomes (PS; ▲), rapamycin (■) or rapamycin-loaded polymersomes (rPS; ▼) using the standard dosage protocol (11 injections, 1 mg/kg rapamycin or equivalent). **a**, tSNE visualization of coreceptor CD40 (**a i**; black), CD80 (**a ii**; purple) and CD86 (**a iii**; forest green) positive B cells, dendritic cells (DCs) and monocyte-and-macrophage-linage cells (M/Ms). **b,** tSNE visualization of major-histocompatibility complex (MHC) II presentation (MHC II+ red; MHC II- blue) on DCs and M/Ms. All data are presented as mean percentage (of DCs or M/Ms) ± SD with *p<0.05; ** p<0.01; *** p<0.001; **** p<0.0001 relative to rPS treatment. Statistical significance was determined by one-way ANOVA with Tukey's multiple comparisons test. AX LN: axillary lymphocenter (deep axillary/axillary/axial and superficial axillary/brachial lymph nodes).

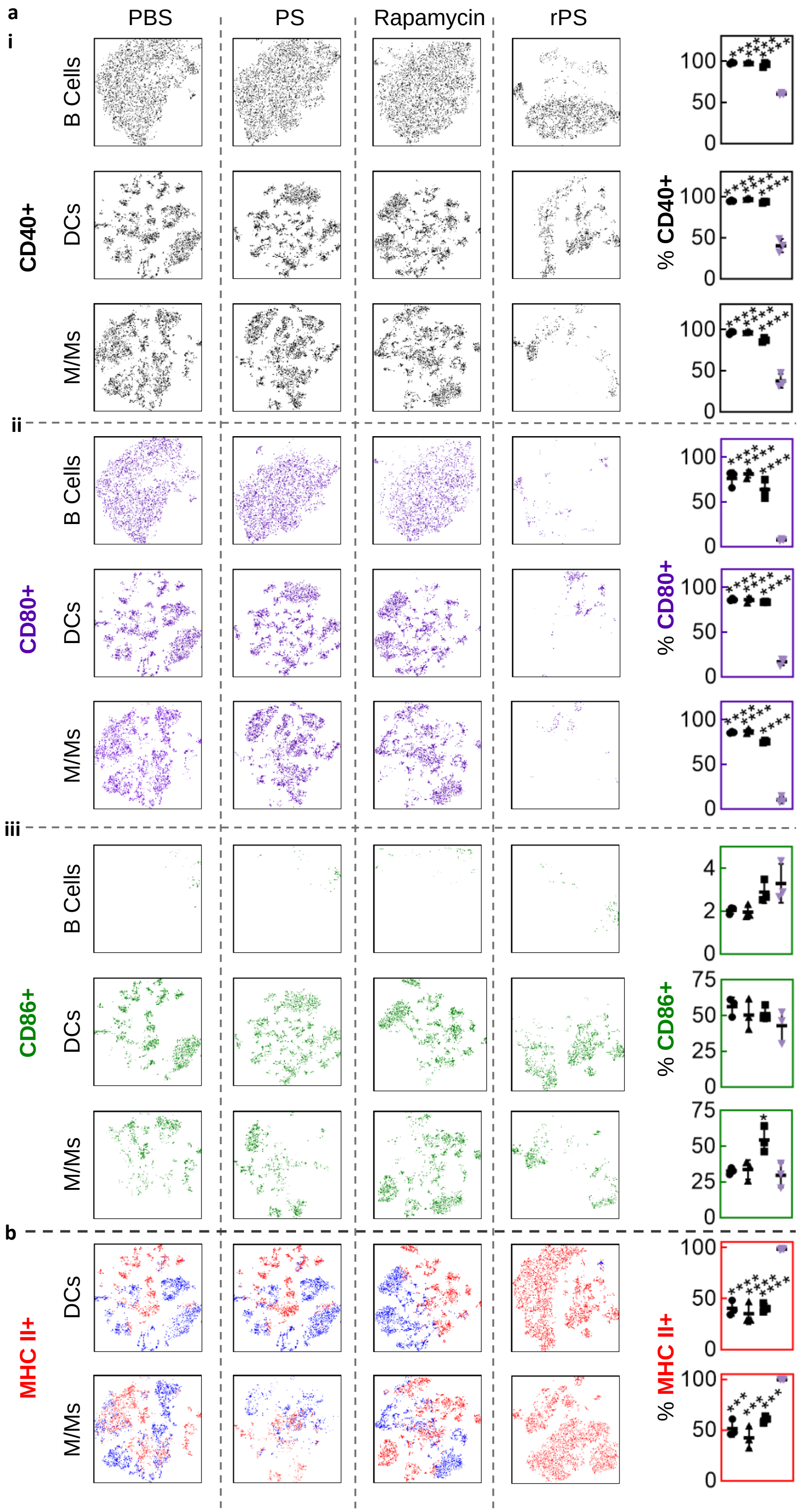

**Fig. S8 | Coreceptor and MHC II expression in IN LN.** Flow cytometry analysis of CD45+ cell populations from mice subcutaneously injected with phosphate suffered saline (PBS; ●), polymersomes (PS; ▲), rapamycin (■) or rapamycin-loaded polymersomes (rPS; ▼) using the standard dosage protocol (11 injections, 1 mg/kg rapamycin or equivalent). **a**, tSNE visualization of coreceptor CD40 (**a i**; black), CD80 (**a ii**; purple) and CD86 (**a iii**; forest green) positive B cells, dendritic cells (DCs) and monocyte-and-macrophage-linage cells (M/Ms). **b,** tSNE visualization of major-histocompatibility complex (MHC) II presentation (MHC II+ red; MHC II- blue) on DCs and M/Ms. All data are presented as mean percentage (of DCs or M/Ms) ± SD with *p<0.05; ** p<0.01; *** p<0.001; **** p<0.0001 relative to rPS treatment. Statistical significance was determined by one-way ANOVA with Tukey's multiple comparisons test. IN LN: subiliac lymphocenter (subiliac/inguinal lymph nodes).

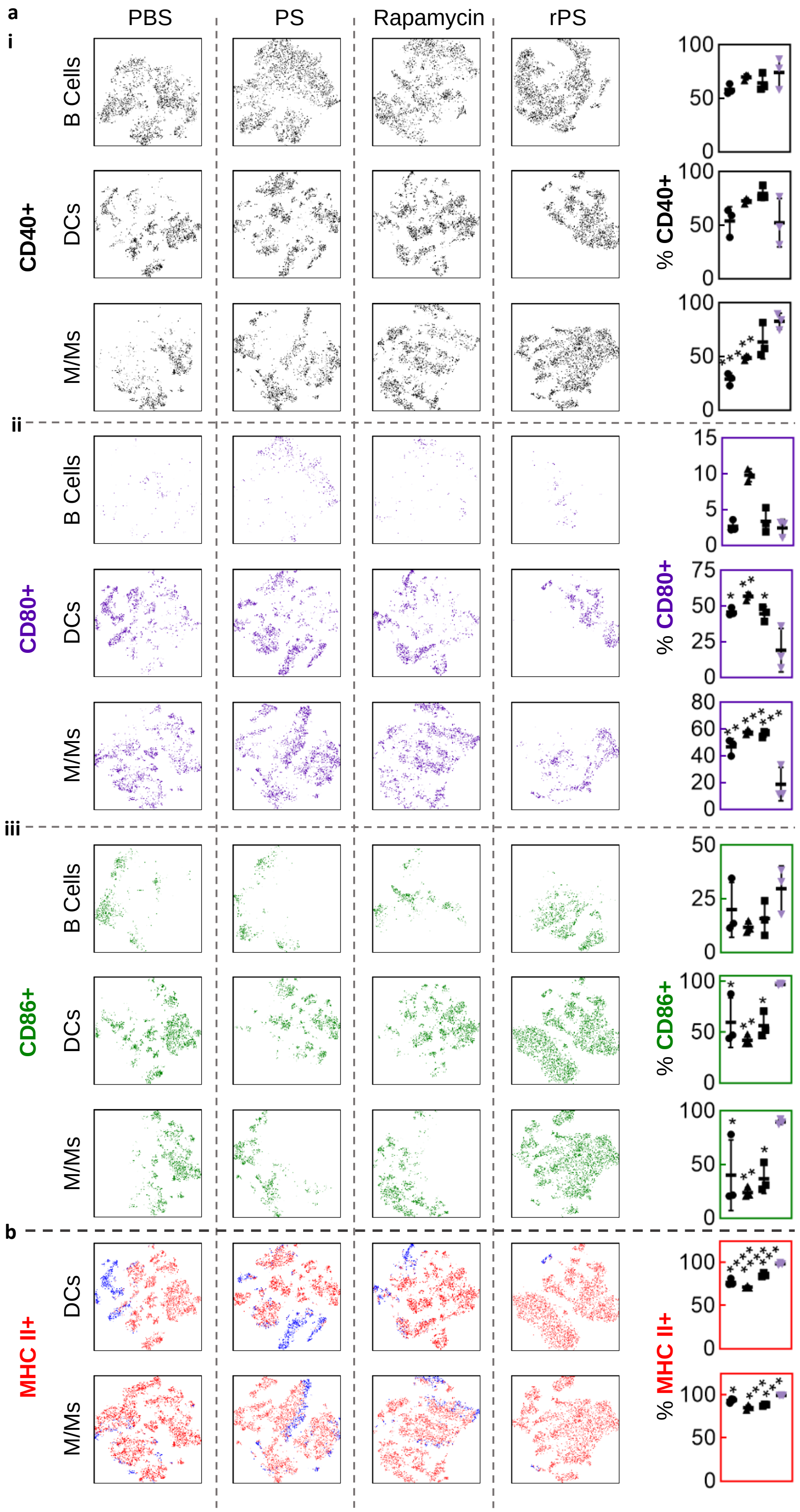

**Fig. S9 | Coreceptor and MHC II expression in the spleen.** Flow cytometry analysis of CD45+ cell populations from mice subcutaneously injected with phosphate suffered saline (PBS; ●), polymersomes (PS; ▲), rapamycin (■) or rapamycin-loaded polymersomes (rPS; ▼) using the standard dosage protocol (11 injections, 1 mg/kg rapamycin or equivalent). **a**, tSNE visualization of coreceptor CD40 (**a i**; black), CD80 (**a ii**; purple) and CD86 (**a iii**; forest green) positive B cells, dendritic cells (DCs) and monocyte-and-macrophage-linage cells (M/Ms). **b,** tSNE visualization of major-histocompatibility complex (MHC) II presentation (MHC II+ red; MHC II- blue) on DCs and M/Ms. All data are presented as mean percentage (of DCs or M/Ms) ± SD with *p<0.05; ** p<0.01; *** p<0.001; **** p<0.0001 relative to rPS treatment. Statistical significance was determined by one-way ANOVA with Tukey's multiple comparisons test.

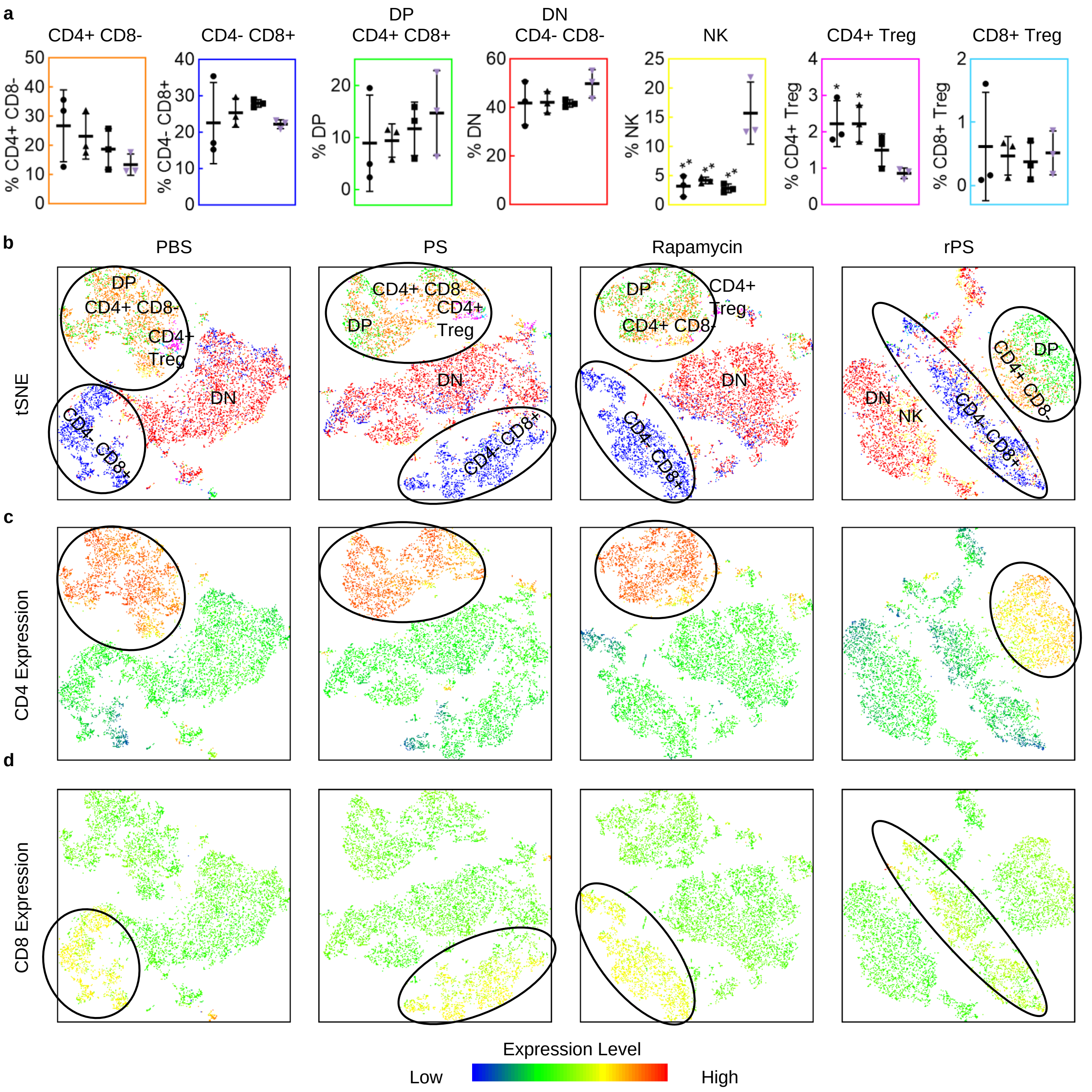

**Fig. S10 | T cell populations in blood.** Flow cytometry analysis of CD3+ T cell populations from mice subcutaneously injected with phosphate suffered saline (PBS; ●), polymersomes (PS; ▲), rapamycin (■) or rapamycin-loaded polymersomes (rPS; ▼) using the standard dosage protocol (11 injections, 1 mg/kg rapamycin or equivalent). **a,** Analysis of CD3+ T cell populations: Percentage of CD3+ T cells (from left to right): CD4+ CD8- (orange), CD4- CD8+ (blue), CD4+ CD8+ double positive (DP; green), CD4- CD8- double negative (DN; red), and natural killer (NK; yellow), CD4+ regulatory (CD4+ Treg; magenta) and CD8+ regulatory (CD8+ Treg; light blue) in blood. All data are presented as mean percentage (of CD3+ cells) ± SD with *p<0.05; ** p<0.01; *** p<0.001; **** p<0.0001 relative to rPS treatment. Statistical significance was determined by one-way ANOVA with Tukey's multiple comparisons test. **b,** tSNE visualization of CD3+ immune cell populations from the liver with color-coded gated overlays of the previously described cell populations: CD4+ CD8- (orange), CD4- CD8+ (blue), CD4+ CD8+ double positive (DP; green), CD4- CD8- double negative (DN; red), and natural killer (NK; yellow), CD4+ regulatory (CD4+ Treg; magenta) and CD8+ regulatory (CD8+ Treg; light blue). **c,d,** tSNE heatmap statistic of CD4 (**c**) and CD8 (**d**) expression. (n=3 mice/group).

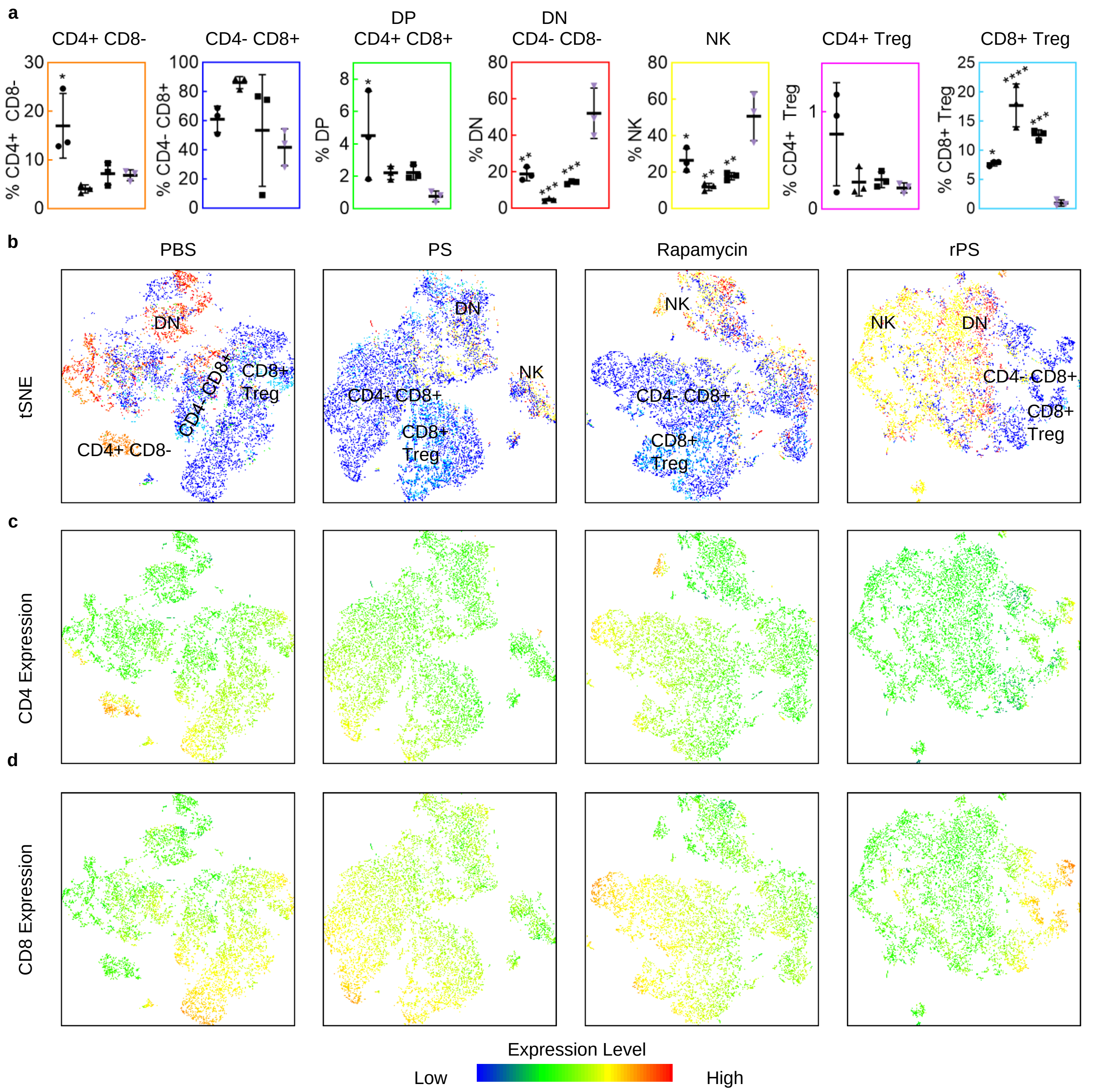

**Fig. S11 | T cell populations in the liver.** Flow cytometry analysis of CD3+ T cell populations from mice subcutaneously injected with phosphate suffered saline (PBS; ●), polymersomes (PS; ▲), rapamycin (■) or rapamycin-loaded polymersomes (rPS; ▼) using the standard dosage protocol (11 injections, 1 mg/kg rapamycin or equivalent). **a,** Analysis of CD3+ T cell populations: Percentage of CD3+ T cells (from left to right): CD4+ CD8- (orange), CD4- CD8+ (blue), CD4+ CD8+ double positive (DP; green), CD4- CD8- double negative (DN; red), and natural killer (NK; yellow), CD4+ regulatory (CD4+ Treg; magenta) and CD8+ regulatory (CD8+ Treg; light blue) in the liver. All data are presented as mean percentage (of CD3+ cells) ± SD with *$p<0.05$; ** $p<0.01$; *** $p<0.001$; **** $p<0.0001$ relative to rPS treatment. Statistical significance was determined by one-way ANOVA with Tukey's multiple comparisons test. **b,** tSNE visualization of CD3+ immune cell populations from the liver with color-coded gated overlays of the previously described cell populations: CD4+ CD8- (orange), CD4- CD8+ (blue), CD4+ CD8+ double positive (DP; green), CD4- CD8- double negative (DN; red), and natural killer (NK; yellow), CD4+ regulatory (CD4+ Treg; magenta) and CD8+ regulatory (CD8+ Treg; light blue). **c,d,** tSNE heatmap statistic of CD4 (**c**) and CD8 (**d**) expression. (n=3 mice/group).

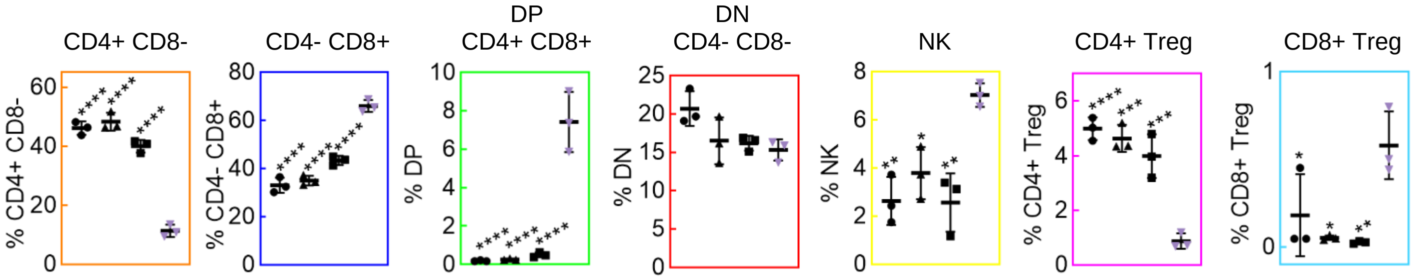

**Fig. S12 | T cell populations in the AX LN.** Flow cytometry analysis of CD3+ T cell populations from mice subcutaneously injected with phosphate suffered saline (PBS; ●), polymersomes (PS; ▲), rapamycin (■) or rapamycin-loaded polymersomes (rPS; ▼) using the standard dosage protocol (11 injections, 1 mg/kg rapamycin or equivalent). **a,** Analysis of CD3+ T cell populations: Percentage of CD3+ T cells (from left to right): CD4+ CD8- (orange), CD4- CD8+ (blue), CD4+ CD8+ double positive (DP; green), CD4- CD8- double negative (DN; red), and natural killer (NK; yellow), CD4+ regulatory (CD4+ Treg; magenta) and CD8+ regulatory (CD8+ Treg; light blue) in the liver. All data are presented as mean percentage (of CD3+ cells) ± SD with *p<0.05; ** p<0.01; *** p<0.001; **** p<0.0001 relative to rPS treatment. Statistical significance was determined by one-way ANOVA with Tukey's multiple comparisons test. (n=3 mice/group). AX LN: axillary lymphocenter (deep axillary/axillary/axial and superficial axillary/brachial lymph nodes).

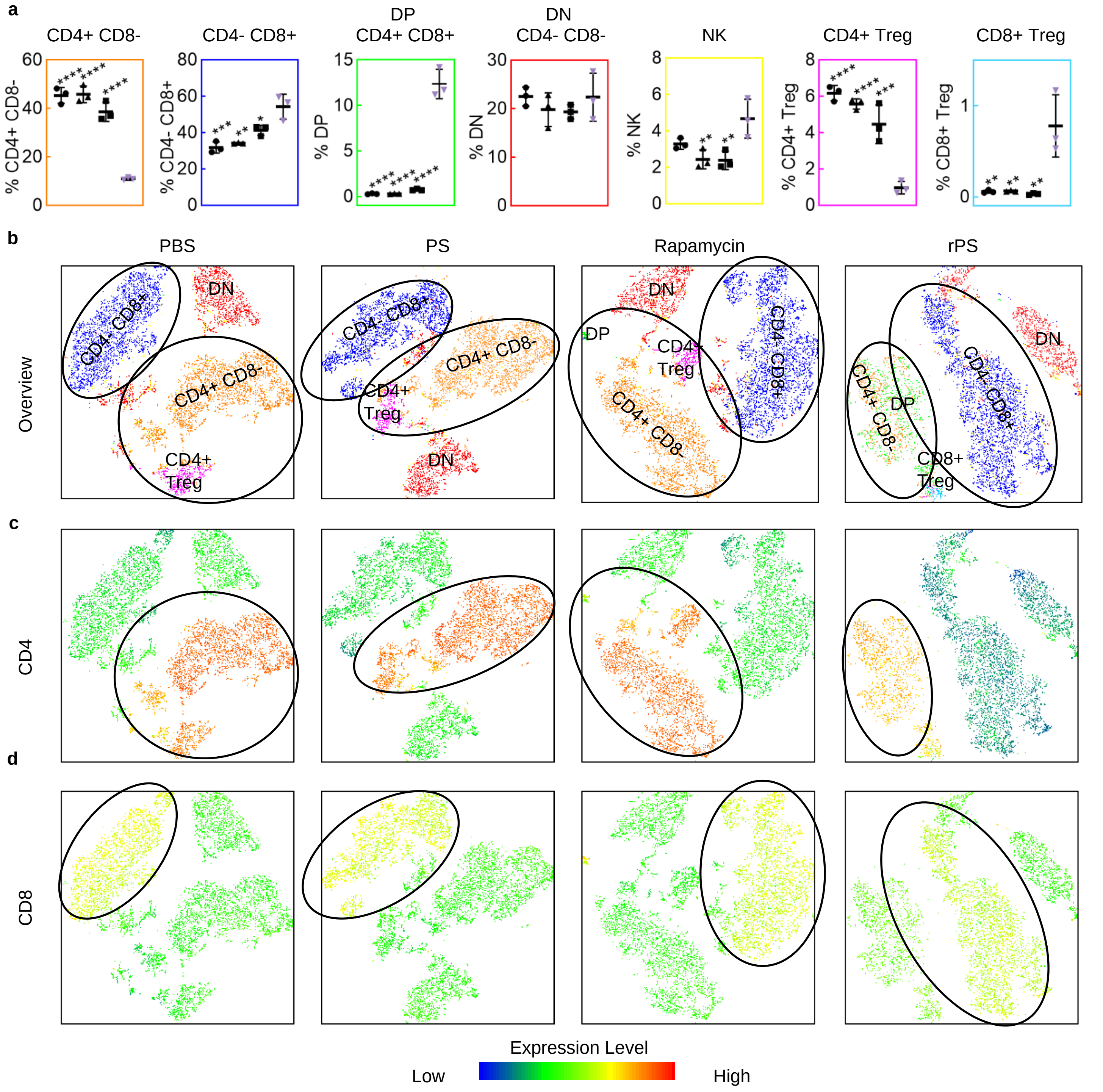

**Fig. S13 | T cell populations in the IN LN.** Flow cytometry analysis of CD3+ T cell populations from mice subcutaneously injected with phosphate suffered saline (PBS; ●), polymersomes (PS; ▲), rapamycin (■) or rapamycin-loaded polymersomes (rPS; ▼) using the standard dosage protocol (11 injections, 1 mg/kg rapamycin or equivalent). **a,** Analysis of CD3+ T cell populations: Percentage of CD3+ T cells (from left to right): CD4+ CD8- (orange), CD4- CD8+ (blue), CD4+ CD8+ double positive (DP; green), CD4- CD8- double negative (DN; red), and natural killer (NK; yellow), CD4+ regulatory (CD4+ Treg; magenta) and CD8+ regulatory (CD8+ Treg; light blue) in the liver. All data are presented as mean percentage (of CD3+ cells) ± SD with *p<0.05; ** p<0.01; *** p<0.001; **** p<0.0001 relative to rPS treatment. Statistical significance was determined by one-way ANOVA with Tukey's multiple comparisons test. **b,** tSNE visualization of CD3+ immune cell populations from the subiliac lymphocenter (subiliac/inguinal lymph nodes; IN LN) with color-coded gated overlays of the previously described cell populations: CD4+ CD8- (orange), CD4- CD8+ (blue), CD4+ CD8+ double positive (DP; green), CD4- CD8- double negative (DN; red), and natural killer (NK; yellow), CD4+ regulatory (CD4+ Treg; magenta) and CD8+ regulatory (CD8+ Treg; light blue). **c,d,** tSNE heatmap statistic of CD4 (**c**) and CD8 (**d**) expression. (n=3 mice/group).

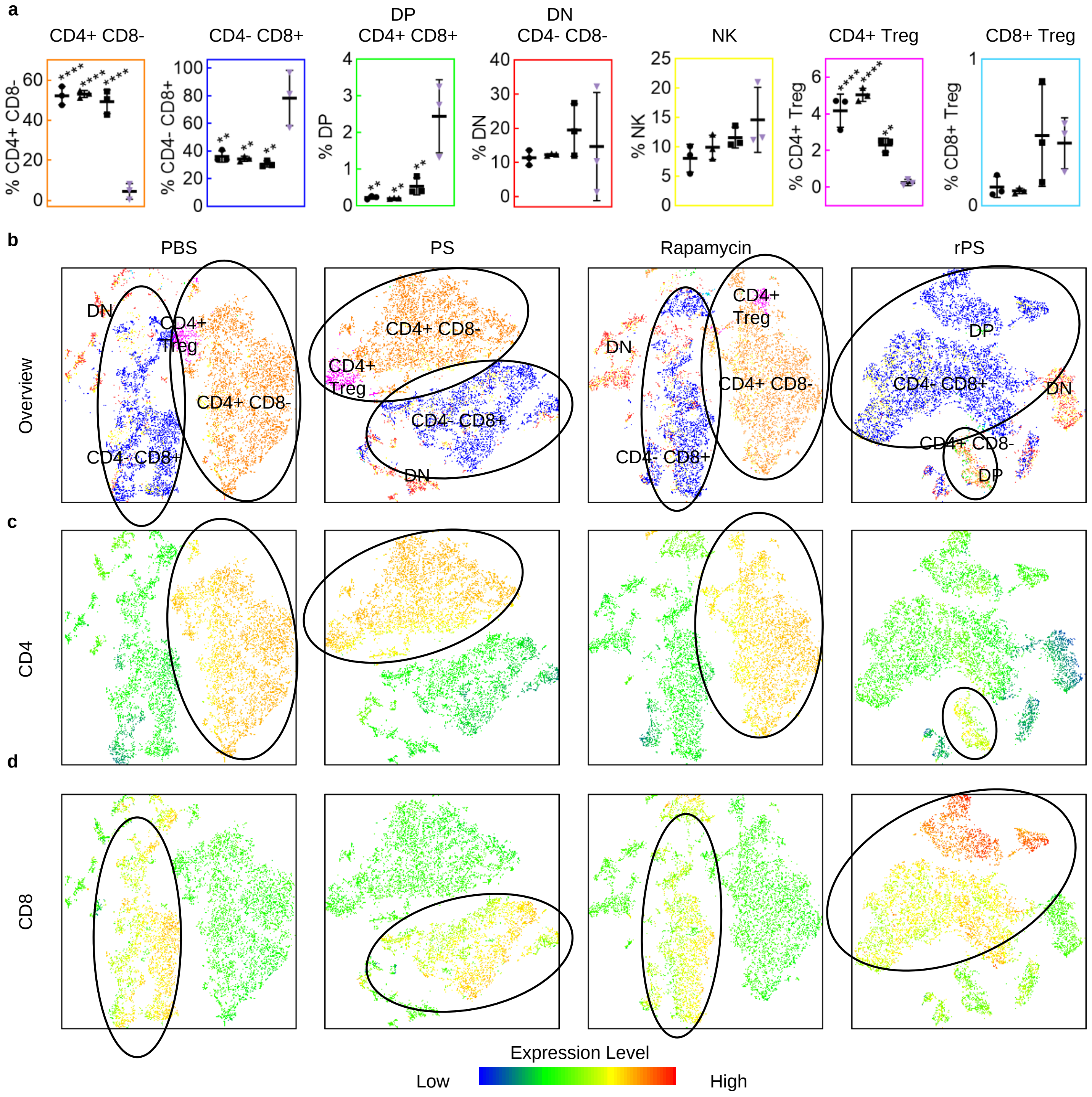

**Fig. S14 | T cell populations in the spleen.** Flow cytometry analysis of CD3+ T cell populations from mice subcutaneously injected with phosphate suffered saline (PBS; ●), polymersomes (PS; ▲), rapamycin (■) or rapamycin-loaded polymersomes (rPS; ▼) using the standard dosage protocol (11 injections, 1 mg/kg rapamycin or equivalent). **a,** Analysis of CD3+ T cell populations: Percentage of CD3+ T cells (from left to right): CD4+ CD8- (orange), CD4- CD8+ (blue), CD4+ CD8+ double positive (DP; green), CD4- CD8- double negative (DN; red), and natural killer (NK; yellow), CD4+ regulatory (CD4+ Treg; magenta) and CD8+ regulatory (CD8+ Treg; light blue) in the liver. All data are presented as mean percentage (of CD3+ cells) ± SD with *p<0.05; ** p<0.01; *** p<0.001; **** p<0.0001 relative to rPS treatment. Statistical significance was determined by one-way ANOVA with Tukey's multiple comparisons test. **b,** tSNE visualization of CD3+ immune cell populations from the spleen with color-coded gated overlays of the previously described cell populations: CD4+ CD8- (orange), CD4- CD8+ (blue), CD4+ CD8+ double positive (DP; green), CD4- CD8- double negative (DN; red), and natural killer (NK; yellow), CD4+ regulatory (CD4+ Treg; magenta) and CD8+ regulatory (CD8+ Treg; light blue). **c,d,** tSNE heatmap statistic of CD4 (**c**) and CD8 (**d**) expression. (n=3 mice/group).

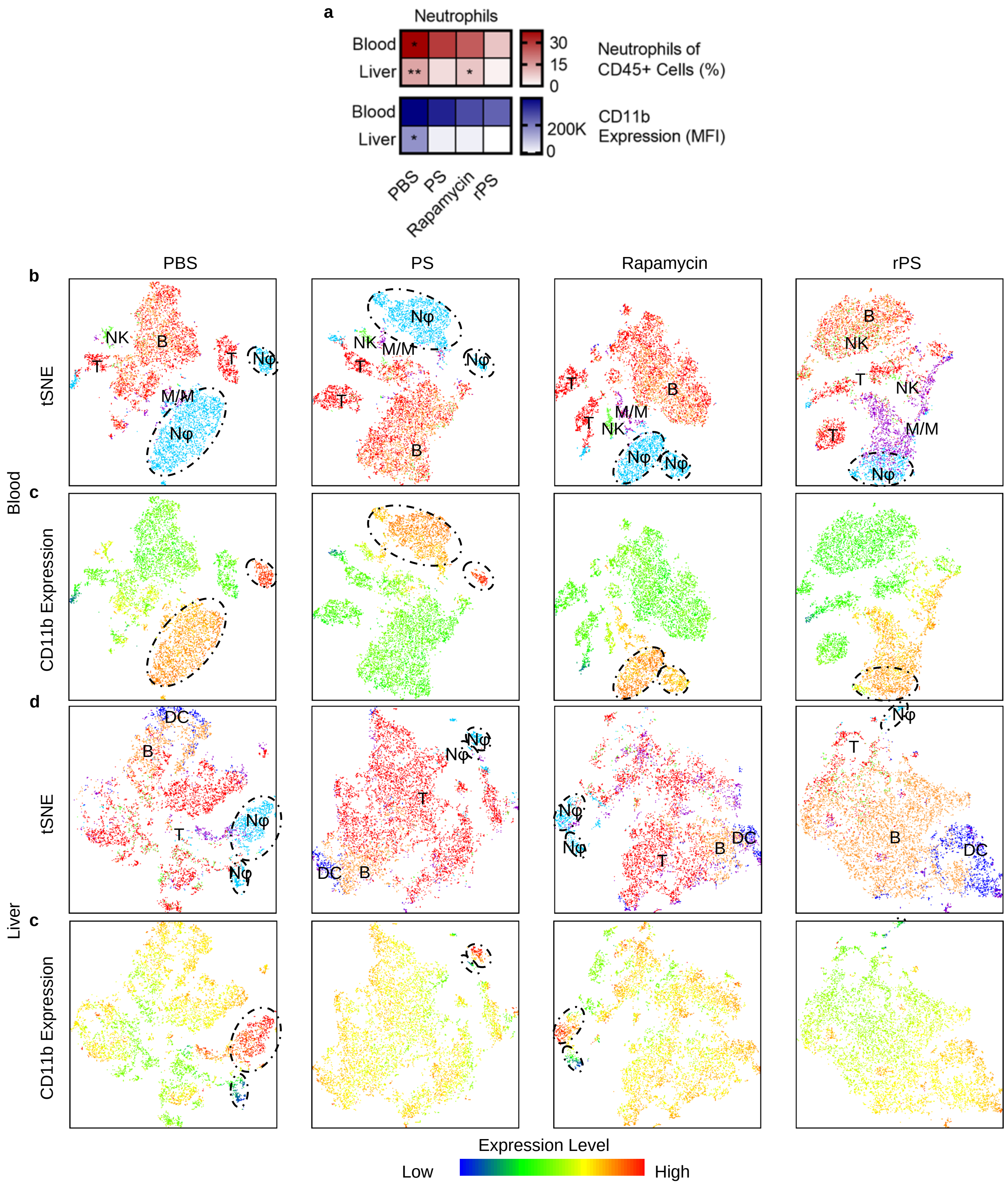

**Fig. S15 | Neutrophil populations in the blood and liver.** Flow cytometry analysis of CD45+ cells from mice subcutaneously injected with phosphate suffered saline (PBS), polymersomes (PS), rapamycin or rapamycin-loaded polymersomes (rPS) using the standard dosage protocol (11 injections, 1 mg/kg rapamycin or equivalent). **a,** Top: Percentage of neutrophils of CD45+ cells in blood and liver. All data are presented as mean percentage (of CD45+ cells) ± SD with *p<0.05; ** p<0.01 relative to rPS treatment. Bottom: CD11b expression by neutrophils in the blood and liver. All data are presented as mean median fluorescent intensity (MFI) ± SD with *p<0.05 relative to rPS treatment. **b,d,** tSNE visualization of CD45+ cells from blood (**b**) and liver (**d**) with color-coded gated overlays of specific cell populations: B cells (B; orange), dendritic cells (DC; blue), monocyte-and-macrophage-linage cells (M/M; purple), neutrophils (Nφ; light blue), and T cells (T; red). **c,e,** tSNE heatmap statistic of CD11b for blood (**c**) and liver (**e**). Dashed-line ovals indicate Nφ populations.

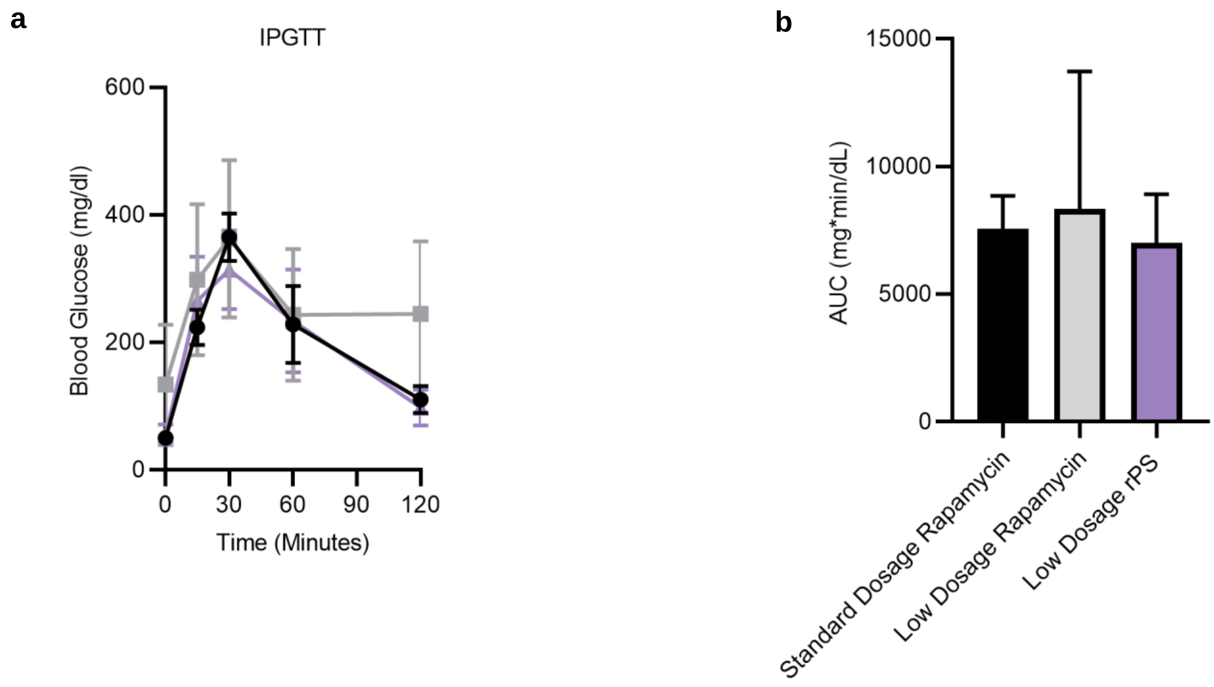

**Fig. S16 | Intraperitoneal glucose tolerance test (IPGTT). a,** Blood glucose concentration over time after intraperitoneal glucose challenge. **b,** Area under the curve from IPGTT. (n = 5-7 mice/group). All data is presented as mean ± SD.

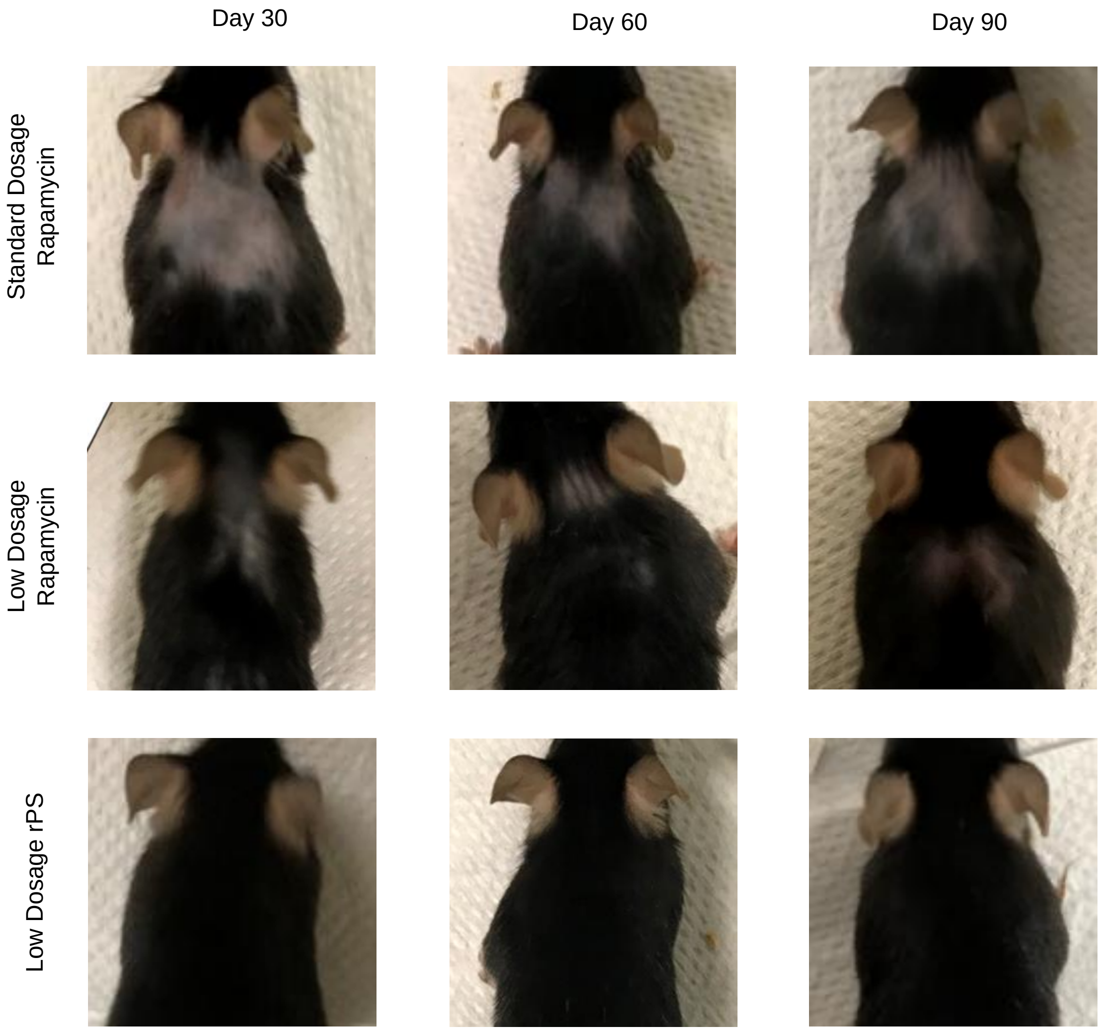

**Fig. S17 | rPS reduce injection site alopecia associated with rapamycin.** (n = 5-7 mice/group).

**Table S1 | Single-cell RNA sequencing analysis workflow**

| Workflow | Program | Command Line |
|---|---|---|
| Quality Control | FastQC v0.11.5 | fastqc <input_path_to/untrimmed.fq.gz> |
| Trimming and Filtering | Trimmomatic v0.39 | java -jar ./Trimmomatic-0.39/trimmomatic-0.39.jar SE -threads 16 -phred33 <input_path_to/untrimmed.fq.gz> <output_path_to/trimmed.fq.gz> ILLUMINACLIP:TruSeq3-SE.fa:2:30:10 LEADING:30 TRAILING:30 MINLEN:36 |
| Alignment | STAR v2.6.0a | STAR --genomeDir ../star_index/ --readFilesCommand zcat –readFilesIn <input_path_to/trimmed.fq.gz>--outFilterType BySJout --runThreadN 16 --outFilterMultimapNmax 100 --alignSJoverhangMin 8 --alignSJDBoverhangMin 1 --outFilterMismatchNoverLmax 0.05 --alignIntronMin 20 --alignIntronMax<br><br>1000000 --alignMatesGapMax 1000000 --outSAMattributes NH HI NM MD --outSAMstrandField intronMotif --outSAMmapqUnique 60 --outSAMtype BAM SortedByCoordinate --outReadsUnmapped<br>Fastx --limitBAMsortRAM 30000000000 --outFileNamePrefix <output_path_to/sorted_bam> |
| Quantification and Differential Expression Analysis | Cufflinks v2.2.1 | cuffdiff -L PBS,PS,R,RPS -o <path_to/output_dir/> -p 16 -b ../genome_fasta/GRCm38.primary_assembly.genome.fa –u ../GTF/gencode.vM24.primary_assembly.annotation.gtf<br>treatment_1_sample_1.bam,treatment_1_sample_2.bam,treatment_1_sample_3.bam<br>treatment_2_sample_1.bam,treatment_2_sample_2.bam,treatment_2_sample_3.bam<br>treatment_3_sample_1.bam,treatment_3_sample_2.bam,treatment_3_sample_3.bam<br>treatment_4_sample_1.bam,treatment_4_sample_2.bam,treatment_4_sample_3.bam |

**Table S2 | Single-cell RNA sequencing raw data**

| **Spleen Macrophages** | | | **PS** | | **Rapamycin** | | **rPS** | | **Rapamycin vs rPS** | |
|---|---|---|---|---|---|---|---|---|---|---|
| | **Ensembl Code** | **Geneset Name** | **Fold Change** | **P Value** | **Fold Change** | **P Value** | **Fold Change** | **P Value** | **Fold Change** | **P Value** |
| 2279 | ENSMUSG00000020053 | IGF1 | -6.39305 | 0.01815 | -9.10778 | 0.00425 | -6.03822 | 0.02255 | 3.06956 | 0.02915 |
| 10384 | ENSMUSG00000036561 | PPP6R2 | -5.69947 | 0.011 | -24 | 0.00005 | -5.5058 | 0.01425 | 24 | 0.00005 |
| 5190 | ENSMUSG00000025786 | ZDHHC3 | -5.15342 | 0.02015 | -8.14259 | 0.03895 | -4.23154 | 0.04065 | 3.91105 | 0.0462 |
| 2460 | ENSMUSG00000020346 | MGAT1 | -2.54064 | 0.0472 | 5.0747 | 0.0171 | -3.04016 | 0.01425 | -8.11486 | 0.001 |
| **Spleen CD4+ Regulatory T Cells** | | | **PS** | | **Rapamycin** | | **rPS** | | **Rapamycin vs rPS** | |
| | **Ensembl Code** | **Geneset Name** | **Fold Change** | **P Value** | **Fold Change** | **P Value** | **Fold Change** | **P Value** | **Fold Change** | **P Value** |
| 573 | ENSMUSG00000003379 | CD79A | -2.52198 | 0.0419 | -24 | 0.00005 | -4.38266 | 0.00455 | 24 | 0.00005 |
| 4444 | ENSMUSG00000024353 | MZB1 | -2.06991 | 0.0297 | -24 | 0.00005 | -2.50151 | 0.0115 | 24 | 0.00005 |
| 947 | ENSMUSG00000006134 | CRKL | -0.689499 | 0.4598 | 7.38707 | 0.02155 | -1.97122 | 0.04935 | -9.3583 | 0.00525 |
| 12235 | ENSMUSG00000041417 | PIK3R1 | 1.96275 | 0.0626 | -12 | 0.00005 | 3.12528 | 0.07685 | 12 | 0.00005 |
| **Liver Macrophages** | | | **PS** | | **Rapamycin** | | **rPS** | | **Rapamycin vs rPS** | |
| | **Ensembl Code** | **Geneset Name** | **Fold Change** | **P Value** | **Fold Change** | **P Value** | **Fold Change** | **P Value** | **Fold Change** | **P Value** |
| 1331 | ENSMUSG00000010651 | ACAA1 | -2.44129 | 0.0089 | -3.83338 | 0.0077 | -1.23789 | 0.03675 | 2.5955 | 0.0395 |
